# Systematic evaluation of protein-small molecule hybrids on the yeast surface

**DOI:** 10.1101/2023.05.12.540568

**Authors:** Manjie Huang, Marina Rueda-Garcia, Abbigael Harthorn, Benjamin J. Hackel, James A. Van Deventer

## Abstract

Protein-small molecule hybrids are structures that have the potential to combine the inhibitory properties of small molecules and the specificities of binding proteins. However, achieving such synergies is a substantial engineering challenge, with fundamental principles yet to be elucidated. Recent work has demonstrated the power of yeast display-based discovery of hybrids using a combination of fibronectin binding domains and thiol-mediated conjugations to introduce small molecule warheads. Here, we systematically study the effects of expanding the chemical diversity of these hybrids on the yeast surface, investigating a combinatorial set of fibronectins, noncanonical amino acid (ncAA) substitutions, and small molecule pharmacophores. Our results show that previously discovered thiol-fibronectin hybrids are generally tolerant of a range of ncAA substitutions and retain binding to carbonic anhydrases following click chemistry-mediated assembly of hybrids with diverse linker structures. Most surprisingly, we identified several cases where replacement of a potent acetazolamide warhead with a substantially weaker benzenesulfonamide warhead still resulted in the assembly of functional hybrids. In addition to these unexpected findings, we expanded the throughput of our system by validating a 96-well plate-based format to produce yeast-displayed hybrid conjugates in parallel. These efficient explorations of hybrid chemical diversity demonstrate that there are abundant opportunities to expand the functions of protein-small molecule hybrids and elucidate principles that dictate their efficient discovery and design.

## Introduction

The precise disruption of individual protein functions is a challenging task, even with the rapidly expanding tools of modern ligand and drug discovery. Clinically approved inhibitors are only available for 3% of the human proteome, only 7% have ligands/inhibitors that are available but not clinically approved, and for more than half of the proteins of the proteome there are not yet any specific inhibitors available.^1^ Although an increasing number of proteins are now considered druggable, specific targeting of individual members of closely related protein families, such as enzymes,^2^ ion channels,^3^ and G protein-coupled receptors (GPCRs),^4–7^ still pose challenging inhibitor discovery problems. Current tools available for specifically interfering with difficult targets are mostly either proteins (antibodies, peptides, or other protein-based binding agents) or small molecules. While engineering highly specific protein binding agents for a given target is becoming more routine, protein-based inhibitors of enzymes remain challenging to discover and engineer.^8–13^ Small molecule drugs can access active sites of target proteins and inhibit their functions efficiently,^14^ but it is difficult to find molecules exhibiting single-target specificity due to the small available surface area for recognition in typical small molecules^15^.

Since proteins and small molecules have complementary features, one strategy for simultaneously leveraging both sets of properties is to prepare “hybrids” that include both protein and small molecule components. Prior efforts in the area have established important proofs-of-concept for the feasibility of generating hybrids with desirable specificities and inhibitory properties. This includes rationally designed hybrid conjugates linking kinase inhibitors to mini-protein scaffolds^16^ or fibronectins^17^ to enhance kinase inhibitor specificity, and conjugation of carbonic anhydrase inhibitors to peptides to generate nanomolar inhibitors.^18, 19^ Recently, Lewis and Harthorn *et al.* constructed combinatorial fibronectin-acetazolamide hybrid libraries and isolated several isoform-selective human carbonic anhydrase (hCA) inhibitors.^20^ These studies highlight the promise of protein-small molecule hybrids and demonstrate the feasibility of high-throughput discovery of such structures. However, systematic investigations of hybrid structure and function remain limited, particularly with respect to the chemical components of hybrids (conjugated amino acids, conjugation chemistries, and warheads). There are substantial opportunities to exploit synergies between protein and small molecule functionality during hybrid discovery and design.

In this study, we established workflows for preparing chemically diverse sets of hybrids on the yeast surface and systematically investigated the properties of the resulting structures (**Figure 1**). We started with a set of previously reported human-carbonic-anhydrase(hCA)-targeting hybrids based on the fibronectin protein scaffold, thiol conjugation sites, and a series of maleimide-PEG-warheads (**Figure 1A**).^20^ Here, we used noncanonical amino acid (ncAA) substitutions and copper-catalyzed azide-alkyne cycloaddition (CuAAC) click chemistry to combinatorially vary the attachment site and hCA-targeting warhead.^21–24^ To increase the throughput of hybrid generation, we also established a 96-well plate-based CuAAC protocol that facilitates rapid assembly and evaluation of hybrid construction and properties in parallel. These investigations allowed us to identify hybrids that retain hCA binding and inhibition while differing substantially from the previously described “parent” hybrids in terms of conjugated amino acid, conjugation chemistry, and even the CA-targeting warhead (with some loss of isoform selectivity noted). Our findings demonstrate the power afforded by the efficient preparation and evaluation of chemically diverse hybrids and reveal opportunities for exploring hybrid “chemical space” to advance the discovery of potent, specific inhibitors.

**Figure 1.**
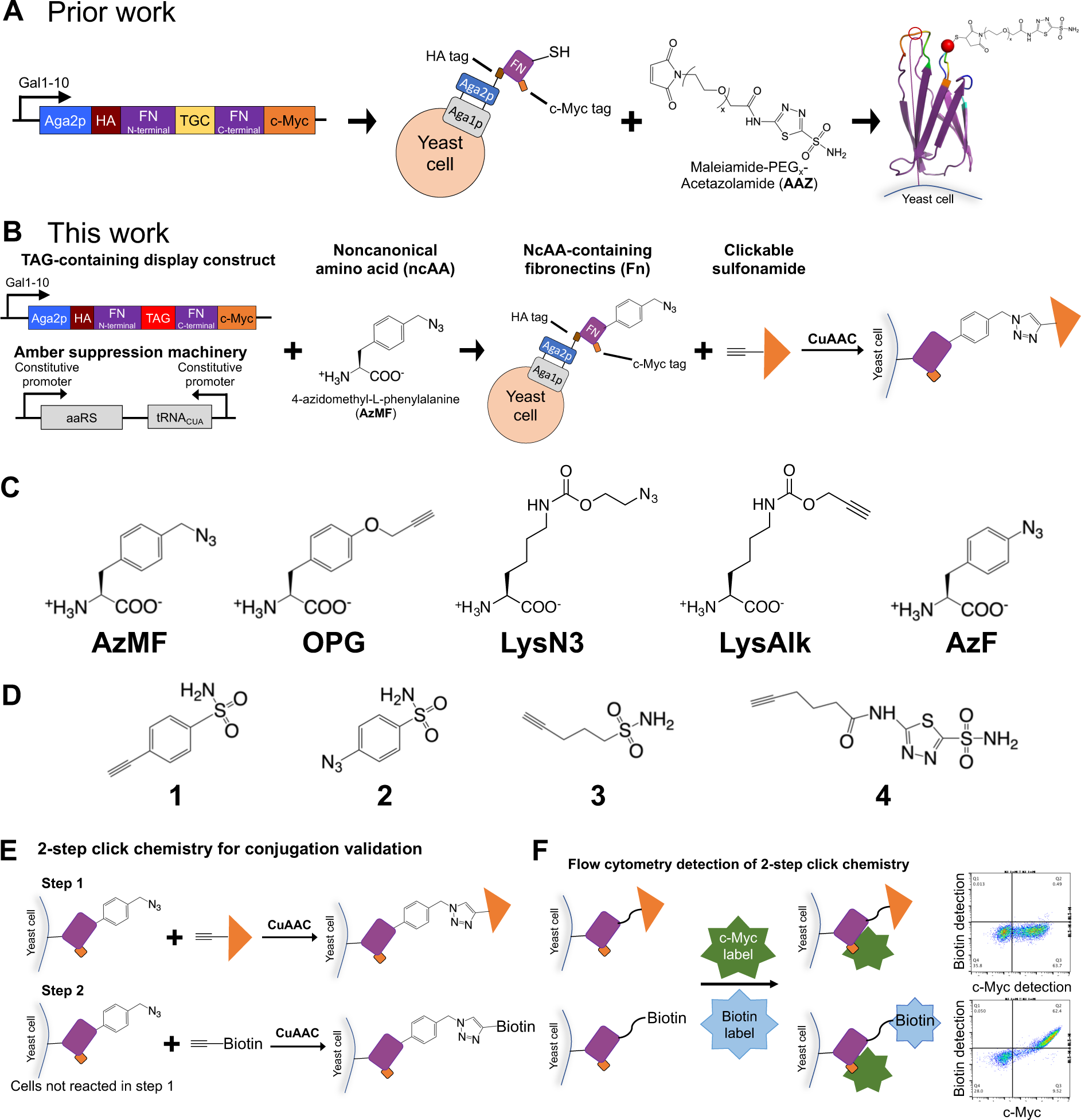
Construction of fibronectin (Fn)-small molecule hybrids using noncanonical amino acids and click chemistry. **A)** Fibronectin-acetazolamide hybrids from prior work. A cysteine positioned at either site 28 (filled red sphere) or site 80 (open red circle) is used for thiol-maleimide chemistry to conjugate PEG-acetazolamide warheads to proteins. **B)** In the present work, the cysteine codon is mutated to a TAG codon so that clickable noncanonical amino acids (ncAAs) can be incorporated at the site by using stop codon suppression machinery (adding a 21^st^ amino acid to these constructs). Small molecule warheads are then conjugated to the ncAA with copper-catalyzed azide-alkyne cycloaddition (CuAAC) chemistry. **C)** Clickable ncAAs used in this work: *p*-azidomethyl-L-phenylalanine (AzMF), *p*-propargyloxyphenylalanine (OPG), *N*-ε-((2-Azidoethoxy)carbonyl)-L-lysine (LysN3), *N*-ε-propargyloxycarbonyl-L-lysine (LysAlk), and *p*-azido-L-phenylalanine (AzF). **D)** Clickable sulfonamides used in this work: 4-ethynyl-benzenesulfonamide (**1**), 4-azidobenzenesulfonamide (**2**), pent-4-yne-1-sulfonamide (**3**), and *N*-(5-sulfamoyl-1,3,4-thiadiazol-2-yl)hex-5-ynamide (**4**). **E)** 2-step click chemistry facilitates validation of click chemistry conjugation via flow cytometry. In the first step, a CuAAC reaction with the cells displaying clickable Fns is performed with a clickable small molecule. Following washing to remove excess reactants from the first reaction, the cells are then clicked with biotin probes in Step 2 to enable flow cytometric detection of ncAAs unreacted following Step 1 using fluorescent streptavidin. **F)** Fluorescent labeling of cells after 2-step click chemistry. Cells are labeled for full length display with anti-cMyc tag labeling and biotin conjugation with fluorescent streptavidin.

## Results and Discussion

### Construction of ncAA-hybrids using yeast display and stop codon suppression

In prior work, Lewis and Harthorn *et al.* reported the discovery of several fibronectin-acetazolamide (Fn-AAZ) hybrids that exhibit selective inhibition of either human carbonic anhydrase II (hCA-II) or hCA-IX. The AAZ pharmacophore was introduced using thiol-maleimide chemistry at engineered cysteine residues, with the AAZ and maleimide groups separated by PEG linkers of varying lengths (**Figure 1A**).^20^

In this work, we diversified several chemical elements of six of the previously discovered hybrids (**Table 1**) with the use of multiple ncAA substitutions and pharmacophores (**Figure 1B**). Genes encoding the fibronectin portion of these hybrids were placed within the pCT40 yeast display vector and mutated to replace the codon encoding the cysteine with a TAG codon. This enabled us to encode ncAAs in response to the TAG codon and display the resulting constructs on yeast using our ncAA-compatible yeast display platform (All orthogonal translation systems used to encode ncAAs in proteins in this work have been previously described).^21, 22^ We used a total of 5 different clickable ncAAs: the four amino acids *p*-azidomethyl-L-phenylalanine (AzMF), *p*-propargyloxyphenylalanine (OPG), *N*-ε-((2-Azidoethoxy)carbonyl)-L-lysine (LysN3), and *N*-ε-propargyloxycarbonyl-L-lysine (LysAlk) were used during initial surveys of binding activity, and one additional ncAA, *p*-azido-L-phenylalanine (AzF) was added during follow-up characterizations (**Figure 1C**). After inducing yeast to display ncAA-containing Fns, we used CuAAC to conjugate clickable small molecules onto the displayed proteins (**Figure 1B**).^24^ We used 4 different sulfonamides (**Figure 1D**) known to inhibit carbonic anhydrases with various potencies. The clickable compound *N*-(5-sulfamoyl-1,3,4-thiadiazol-2-yl)hex-5-ynamide (**4**, K_i_ ∼10 nM) contains the same acetazolamide (AAZ) pharmacophore used during prior hybrid discovery efforts in Fn-Cys form.^20^ 4-ethynyl-benzenesulfonamide (**1**, K_i_ ∼100-1000 nM) and 4-azidobenzenesulfonamide (**2**, K_i_ ∼100-1000 nM) are clickable aryl sulfonamides with potency reduced compared to the AAZ warhead, while pent-4-yne-1-sulfonamide (**3**, K_i_ >10,000 nM) is an alkyl sulfonamide with further reduced potency in the micromolar range.^25–27^ Since not all of these warheads can be directly detected on the yeast surface, we used our established 2-step click chemistry protocol to evaluate conjugation reactions on yeast surface (**Figure 1E**).^24^ In the first step, cells displaying the fibronectins are subjected to reactions with the sulfonamides and washed. In the second step, cells are then subjected to click chemistry with a biotin probe to detect the presence of ncAAs that did not react with the small molecule. After additional washing, cells are then labeled to detect biotin and full-length proteins and subjected to flow cytometric analysis (**Figure 1F**).^24^ In general, this resulted in reduction of biotin detection to background levels (**Figure S1**), corresponding to estimated extents of reaction of approximately 90% (**Supplementary Table S1**). However, for the case of conjugations with AAZ-alkyne, when we used longer reaction times or higher AAZ concentrations in an attempt to achieve high extents of reaction, we observed that displayed fibronectins lose binding function. Therefore, we settled for lower extent of reactions in AAZ-treated samples in order to ensure the presence of functional displayed hybrids (**Figure S2**, **Supplementary Table S2**). In conclusion, we successfully constructed 48 distinct hybrids on the yeast surface and verified their preparation using flow cytometry.

**Table 1.** Previously reported thiol-Fn hybrid inhibitory constants determined (K_i_) in solution.

| hCA-II specific | hCA-IX specific |
| --- | --- |
| FnII.28.5-PEG<sub>5</sub>-AAZ ($K_i$ = 24 nM) | FnIX.28.2-PEG<sub>2</sub>-AAZ ($K_i$ = 8.7 nM) |
| FnII.28.7-PEG<sub>7</sub>-AAZ ($K_i$ = 4.8 nM) | FnIX.28.5-PEG<sub>5</sub>-AAZ ($K_i$ = 3.4 nM) |
| FnII.80.7-PEG<sub>7</sub>-AAZ ($K_i$ = 88 nM) | FnIX.28.7-PEG<sub>7</sub>-AAZ ($K_i$ = 48 nM) |

### Survey and characterization of hybrids using flow cytometry

Following hybrid construction, we surveyed the binding activities of the Fn-sulfonamide hybrids following incubation of hybrid-displaying cells with either 50 nM hCA-II (His-tag) or hCA-IX (human Fc tag) (**Figure 2A**).

**Figure 2.**
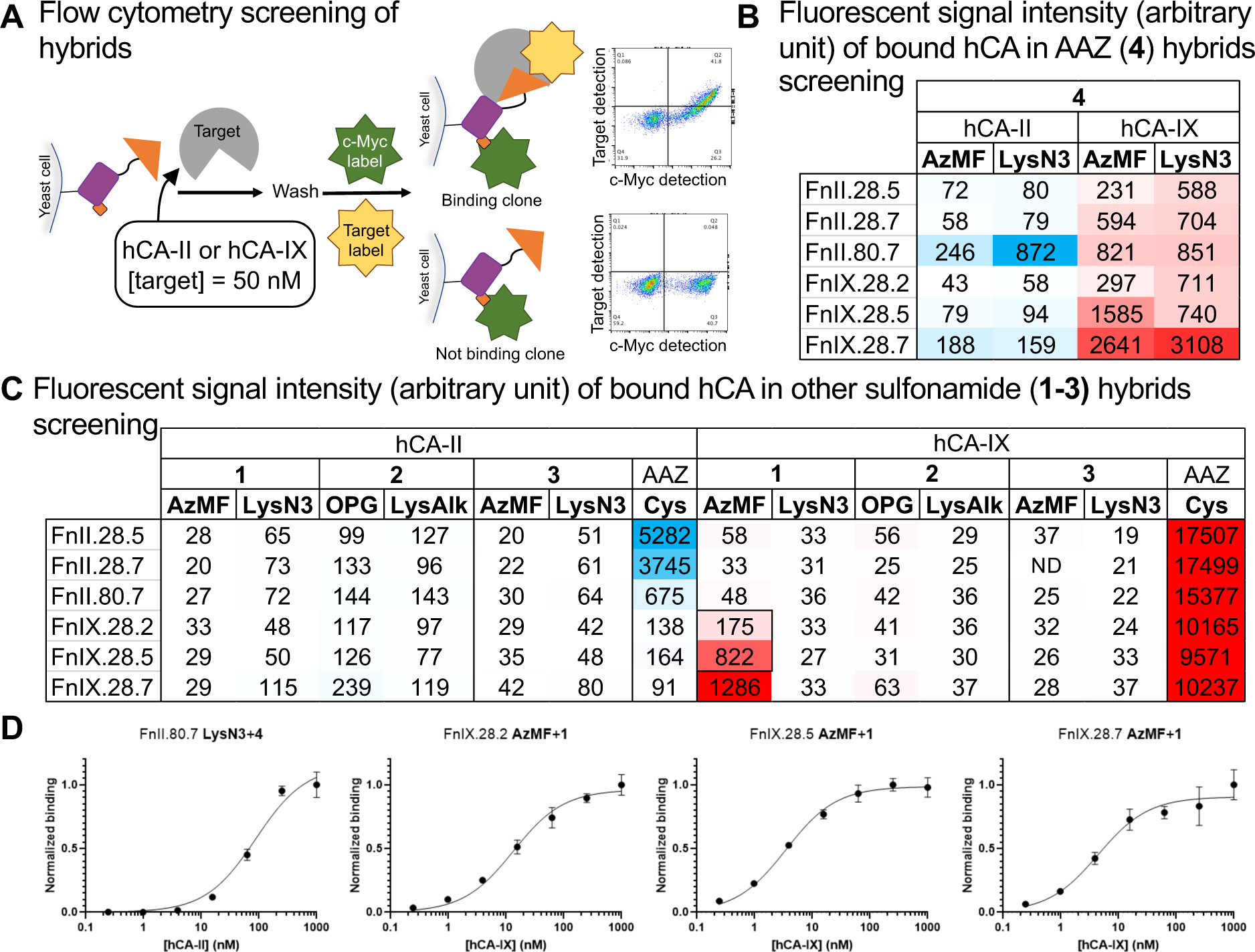
Binding assays with yeast-displayed hybrids reveal multiple hCA-binding, Fn-small molecule hybrids prepared via click chemistry. **A)** Hybrids are evaluated for binding properties using flow cytometry. In initial assays on Fn-small molecule hybrids, cells displaying hybrids were incubated with 50 nM hCA-II or hCA-IX and then labeled for C-terminal c-Myc tag (x-axis) and the bound hCA (y-axis). The hCA-II used contains 6x His tag and is labeled with mouse anti-6x His tag antibodies (3D5) and goat anti mouse IgG – Alexa Fluor™ 488. The hCA-IX used contains human Fc tag and is labeled with goat anti human – Alexa Fluor™ 488. **B and C)** Fluorescent signal level detected from hybrids in the hCA binding screening, corresponding to **Supplementary Figure S3-5**. Numbers are median fluorescent intensity (arbitrary units) of hCA labeling fluorophores on hybrid displaying (c-Myc positive) cells subtracting the background fluorescence from the cells not displaying the construct (c-Myc negative). The arbitrary units of fluorescence detection necessitate relative comparisons. Heat map color intensity scales were configured such that binding hybrids we identified from dot plots are readily apparent. **D)** Exemplary yeast surface titrations to estimate affinities of the hybrids for target hCAs. The full set of titration curves obtained on hybrids are shown in **Supplementary Figure S7**, and the estimated KD values are reported in **Table 2**.

As positive controls, we conjugated maleimide-PEG_7_-acetazolamide to the “parent” cysteine-containing Fns and subjected them to identical binding assays. In the flow cytometry data, we gated on cells displaying full-length proteins (c-Myc positive cells) and evaluated binding behavior in only these subpopulations. Heat maps of median fluorescence intensities enable evaluations of binding activity as a function of Fn clone, ncAA substitution, warhead, and hCA isoform (**Figure 2B, C**). These single-concentration hCA binding assays enabled us to rapidly identify hybrids exhibiting binding activities. Notably, data from these experiments cannot directly be used to evaluate relative binding affinities since these series of ncAA-substituted Fns exhibit variable display levels that are not taken into account when detecting hCA binding (note sample-to-sample changes in c-Myc levels in **Supplementary Figures S3-5**).

Click chemistry-mediated installation of the AAZ warhead (small molecule **4**) resulted in numerous hybrids exhibiting binding to hCA isoforms (**Figure 2B, Supplementary Figure S3**). This is consistent with the use of an AAZ warhead during the initial discovery of the Fn clones used here as well as its relatively high affinity. Interestingly, all six AAZ-modified hybrids exhibited hCA-IX binding using both AzMF and LysN3 as ncAA attachment points, while only two clones (FnII.80.7 and FnIX.28.7) showed detectable hCA-II binding under the conditions tested here. The three clones known to exhibit selectivity for inhibition of hCA-IX in AAZ-thiol-Fn form all bound to hCA-IX in AAZ-azide-Fn form but tended to lack binding to hCA-II, with two clones exhibiting no detectable hCA-II binding and one exhibiting low levels of binding. In contrast to the retention of hCA-IX binding, most of the hybrids previously reported to be selective for hCA-II do not appear to retain hCA-II binding following ncAA substitution and click-mediated AAZ installation except for FnIX.80.7. This hybrid exhibited binding to both hCA-II and -IX with either LysN3 or AzMF substitutions. Interestingly, binding to hCA-IX but not hCA-II was detected with the AAZ-azide-Fn forms of both FnII.28.5 and FnII.28.7. While these molecules exhibited hCA-II preferences in AAZ-thiol-Fn forms, there are important differences that may explain observation of hCA-IX binding in the current context. The “clicked” linkers used in this study are shorter and more rigid than the PEG-based linkers used during the discovery of the isoform-selective clones. The changes in isoform selectivity are consistent with prior work: when PEG-based linkers were shortened, this strongly affected isoform selectivity. Our data presented here raises the possibility that linker rigidity may also play a role in determining isoform selectivity (although further work is needed to more specifically examine the role of linker rigidity to control for changes in length). Overall, hybrids still exhibit binding function following substantial changes in linker structure without changing the warhead, but isoform selectivity and other nuanced hybrid properties are altered when distinct linkers are used.

We also evaluated the binding activities of Fn hybrids prepared with aryl and alkyl sulfonamides (**1**-**3**) known to possess significantly weaker potencies than AAZ (**Figure 2C, Supplementary Figure S4, S5**). While most of these combinations did not result in hybrids exhibiting detectable CA binding activity, the three hCA-IX-targeting clones prepared with AzMF and alkyne sulfonamide **1** exhibited substantial binding to hCA-IX (FnIX.28.7 with AzMF+**1** was also found to bind to hCA-II). These activities are surprising after the drastic changes we made to the hybrid structure from the initial Fn-Cys-AAZ hybrids. The tolerance of a distinct warhead is particularly notable because the aryl sulfonamide was not used during any portion of initial hybrid discovery campaigns and is less potent than the AAZ warhead by at least an order of magnitude. In addition, AzMF clicked with **1** possesses multiple rigid ring structures and a much shorter overall length than Cys modified with maleimide-PEG-AAZ linkers. Interestingly, we did not detect binding activity in any hybrids substituted with OPG and modified with aryl sulfonamide **2**, despite the fact that the warhead is identical to the warhead in **1**. The reversal in polarity of the triazole ring formed during CuAAC and the additional ether group in the side chain of OPG (relative to AzMF) could be less favorable for hCA binding compared to the AzMF + **1** combination. Perhaps unsurprisingly, in no cases did we observe binding when the very weak alkyl sulfonamide warhead **3** was installed, consistent with its potency being several orders of magnitude weaker than AAZ. Overall, these results demonstrate that exploring more combinations of linker and small molecule pharmacophores can reveal unexpected functional hybrids.

We then chose several hybrids that exhibited high hCA binding in our initial binding surveys and subjected them to titrations on the yeast surface to estimate binding affinities (**Figure 2D, Supplementary Figure S6, S7**). From the apparent K_D_s determined here, “clicked” hybrids exhibit single- to triple-digit nM target binding (**Table 2**).

**Table 2.** Yeast surface binding affinities of exemplary hybrids.

| Fibronectin clone | ncAA+small molecule | Target hCA | K<sub>D</sub> (95% CI), nM |
| --- | --- | --- | --- |
| FnII.80.7 | **LysN3+4** | hCA-II | 88 (68 - 110) |
| FnIX.28.2 | **AzMF+1** | hCA-IX | 13 (10 - 17) |
|  | **AzF+1** |  | 10 (8.0 - 14) |
| FnIX.28.5 | **AzMF+1** | hCA-IX | 3.5 (2.9 - 4.2) |
|  | **AzMF+4** |  | 27 (20 - 38) |
|  | **AzF+1** |  | 46 (35 - 61) |
| FnIX.28.7 | **AzMF+1** | hCA-II | 70 (48 - 100) |
|  |  | hCA-IX | 4.4 (3.0 - 6.5) |
|  | **AzMF+4** | hCA-II | 74 (51 - 110) |
|  |  | hCA-IX | 110 (85 - 130) |
|  | **AzF+1** | hCA-IX | 62 (46 - 83) |
|  | **LysN3+4** |  | 170 (140 - 220) |

For comparison, the inhibition constants of the original Fn-AAZ hybrids are shown in **Table 1**.^20^ Although these inhibition constants are from in-solution inhibition assays instead of yeast surface binding titrations, similarities between these values indicate that “clicked” hybrids can achieve comparable potencies against hCAs even after drastic changes in structures. Despite significant changes in molecular design, and without protein evolution, clicked Fn-AAZ hybrids exhibit nM affinity towards their targets. In the case of FnIX.28.5 and FnIX.28.7, construction of hybrids with AzMF + AAZ **4** results in weaker K_D_s to hCA-IX in comparison to the hybrids with AzMF + benzene sulfonamide **1**. A hybrid form of FnIX.28.7 constructed with LysN3 + AAZ **4** exhibited the highest binding signal in initial binding assays, yet the titration data revealed a triple-digit nM affinity to hCA-IX. In contrast to several other clicked hybrids, detection of hCA-IX did not plateau under the conditions tested here. This likely leads to an underestimate of the binding affinity of this hybrid. One possible explanation for this phenomenon is that hCA-IX is known to dimerize at high concentrations,^28^ and the hCA-IX used in this work is fused to human antibody constant region (human Fc), which could lead to further multimerization. This may result in increased antigen binding signal even after all binding sites on hybrids are occupied. The hybrids with the highest apparent affinities on yeast are the hCA-IX targeting clones with AzMF and benzenesulfonamide **1,** exhibiting low double- to single-digit nM level apparent K_D_s. These strong binding affinities confirm that potent hybrids can be generated even starting from relatively weak pharmacophores. Finally, we switched AzMF to a similar ncAA, AzF, which is one carbon shorter than AzMF (**Figure 1C**) and conducted binding titrations. This small change in the ncAA structure changed the affinities of FnIX.28.5-PEG_5_-AAZ and FnIX.28.7-PEG_7_-AAZ for hCA-IX from single-digit nM to double-digit nM values. This suggests that the ncAA side chains in these hybrid structures play an important role in target binding and further confirms that the “chemical details” of hybrids dictate hybrid performance. Overall, these studies demonstrate the feasibility of using ncAA-compatible yeast display as a platform for chemically diversifying existing hybrids to efficiently investigate how different hybrid elements enhance or alter hybrid properties.

### High-throughput fibronectin hybrid generation and screening

The strong performance of multiple new hybrids despite a limited evaluation of molecular space during the initial screening led us to investigate strategies for increasing the throughput of hybrid generation and evaluation. Our “conventional” approach to CuAAC on the yeast surface in microcentrifuge tubes limits experimental throughput (20-30 reactions per experiment; one-by-one handling of samples). To increase the throughput of hybrid construction and screening, we developed a 96-well plate-based click chemistry protocol that allowed us to chemically modify more than 100 samples a day by hand since the fluid handling time is reduced with the use of multichannel pipettes and the lack of need to transfer cells between tubes and plates (**Figure 3A**).

**Figure 3.**
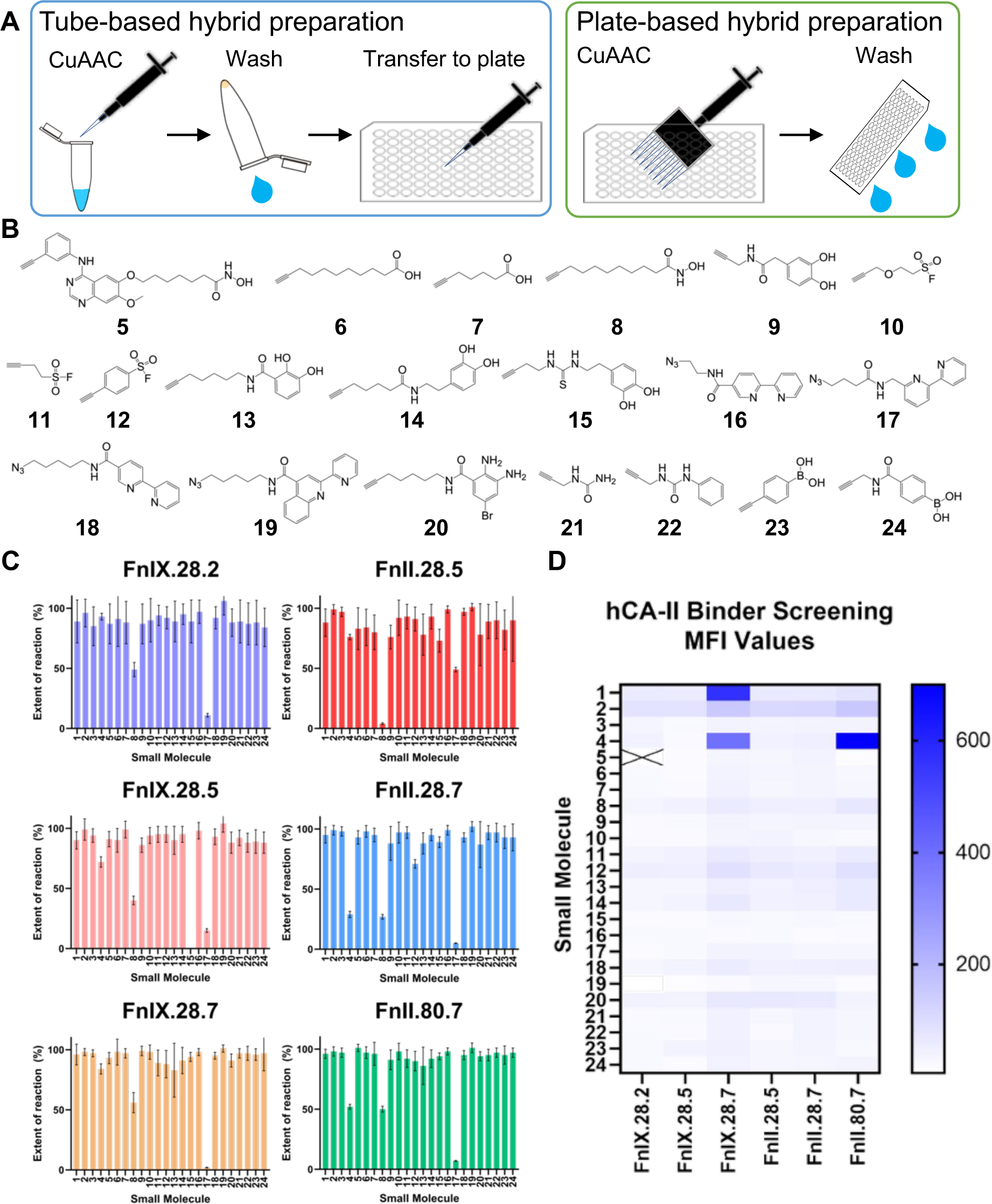
High-throughput hybrid construction and screening. **A)** Comparison between “conventional’ tube-based CuAAC protocol and plate-based CuAAC protocol. Hybrid generation throughput is greatly increased by using multichannel pipette and 96-well plate, and the amount of liquid handling is also reduced. **B)** Additional clickable small molecules used in high-throughput hybrid generation along with small molecule **1**-**4**. Names of the small molecules are listed in Supplementary table S7. **C)** Extent of CuAAC of the 24 small molecules in this screen. **D)** Fluorescent signal level detected from hybrids in the hCA-II binding screening. Numbers are calculated as described in Figure 2. Since the numbers are of arbitrary units of fluorescence detection, the number value is only good for relative comparisons, therefore binding clones are highlighted using an arbitrary intensity scale of the heatmap.

To determine the feasibility of the plate-based approach, we verified that reactions conducted in plates result in indistinguishable outcomes from reactions conducted in tubes, and that no reactant or sample cross contamination between adjacent wells was detected in individual wells of the plate (**Supplementary figure S8,9**). We also tested different combinations of temperature, cell density and small molecule concentrations (**Supplementary figure S10-12**) and verified that CuAAC reactions remain efficient over varied conditions.

To illustrate the utility of this plate-based protocol, we constructed 144 hybrids from 24 clickable small molecules and the six fibronectins described above. After evaluating extent of reaction for each conjugate, we evaluated the resulting conjugates for hCA-II binding. All 24 small molecules we used have functional groups that can target catalytic residues or interact with metal ions commonly found in metalloenzymes (**Figure 3B, Table S7**). From the 2-step characterization of extent of reaction, we determined that most of the hybrids are constructed with greater than 90% extent of reaction (**Figure 3C**). Small molecules **4**, **8**, and **17** all appear to react inefficiently with multiple fibronectin clones. The inefficient reaction of **4** in plates is consistent with tube-based reactions described above. The small molecules **8** and **17** each possess functional groups capable of chelating copper ions and thus hinder CuAAC reactions. However, other molecules that exhibit high reactivity also possess such groups (e.g. **16**, **18**, and **19**); further investigations to better understand why these differences in reactivity occur are merited in future work. In binding assays, we identified 3 hCA-II binding hybrids, including two that were found in the initial tube-based survey (FnII.80.7 with AzMF and **4**, FnIX.28.7 with AzMF and **4**), and one weak binder that was not detected in initial experiments: FnIX.28.7 AzMF conjugated with **1** (**Figure 3B**). In the tube-based assays, this hybrid did not exhibit detectable levels of hCA-II binding, while in the plate-based assays the binding signal is comparable to that of FnII.80.7 with AzMF and **4** (**Figure 3D**); this is lower than the best hCA-II binder, FnII.80.7 with LysN3 and **4** (**Figure 2B**). We attribute the loss of weaker binding signals in these assays to the decay of fluorescent signal over time. We verified that this phenomenon results in position-dependent decreases in binding signal in well plates where the order in which wells are analyzed via flow cytometry affects fluorescence levels (**Supplementary Figure S13**). In future assays this effect can be reduced by using higher affinity primary and secondary detection reagents and by decreasing the per-sample collection time during flow cytometry analysis. Nevertheless, the current implementation represents an effective screening tool. Looking at the full set of hybrids tested here, these proof-of-principle assays did not reveal any hCA-II binding hybrids containing pharmacophores other than aryl sulfonamide **1** or acetazolamide **4**. In any case, this validated plate-based protocol for hybrid construction can be enhanced further with automated liquid handling to explore larger pharmacophore and linker portions of “hybrid space” in future work.

### Solution-based characterizations of ncAA-Fn hybrids

To investigate how observations on the yeast surface transfer over into the solution phase, we prepared FnIX.28.7 AzMF in soluble form. To facilitate expression and purification, we used a previously established yeast protein secretion strategy that supports ncAA incorporation^29^ in which constructs of interest are fused with the human antibody constant region (Fc) (**Figure 4A**).

**Figure 4.**
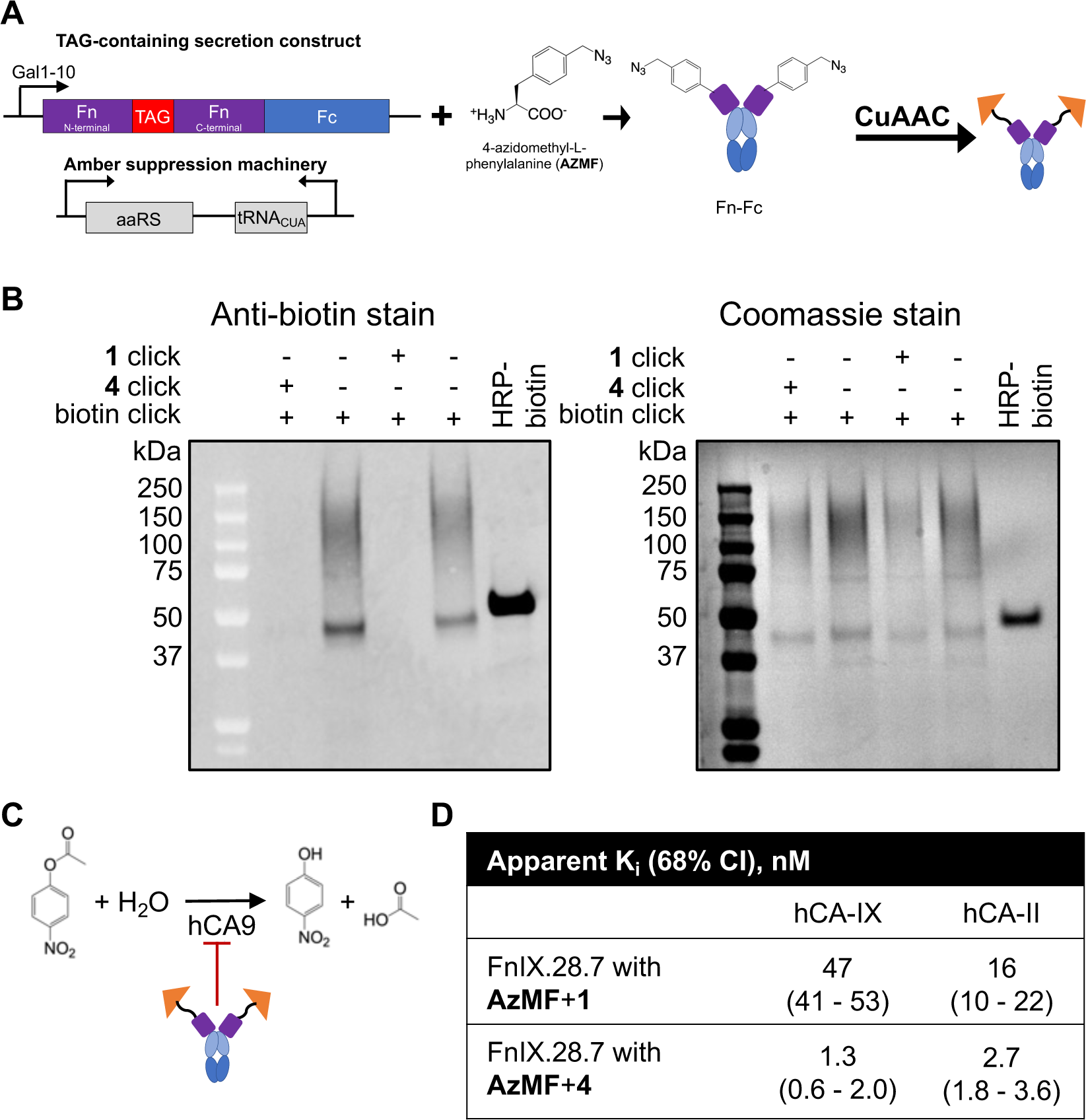
Soluble Fn-sulfonamide hybrids inhibits hCA-II and hCA-IX in solution. **A)** Construction of soluble Fn-sulfonamide hybrid. Fns are fused with human antibody constant region (Fc) and secreted by the yeast after induction. The proteins are then purified and clicked with sulfonamide small molecules to form the soluble Fn-sulfonamide hybrids. **B)** 2-step click chemistry confirms the formation of soluble Fn-sulfonamide hybrids. The Fn-Fcs conjugated with small molecules show no detectable biotin conjugation in the second click chemistry, showing that the click conjugation was successful. **C)** Activity of hCA is measured by the rate of conversion of 4-nitrophenyl acetate into 4-nitrophenol and acetic acid. **D)** The apparent Ki of FnIX.28.7 AzMF conjugated with either benzenesulfonamide (**1**) or acetazolamide (**4**) are calculated in MATLAB. The estimated active hCA-IX concentration is ∼42 nM, and the active hCA-II concentration is ∼23 nM. The nominal hCA concentration is 100 nM for both hCA-II and IX.

We successfully purified FnIX.28.7 AzMF using protein A affinity chromatography (**Supplementary Figure S14**). We used two-step click chemistry in solution to verify the modification of the construct with desired small molecules as follows: we subjected the construct to CuAAC with either small molecule **1** or **4**, removed excess reagents via buffer exchange, and then subjected the construct to click chemistry with a biotin probe.^24^ Subsequent SDS-PAGE and Western blot experiments show that the purified fibronectin can be clicked with biotin, but following CuAAC with either small molecule the putative hybrids cannot, strongly suggesting the successful construction of the desired hybrids in solution (**Figure 4B**).

We investigated the properties of soluble hybrids with a series of inhibition assays (**Figure 4C**, **Supplementary Table S8**). In the inhibition assays we confirmed that the hybrids inhibit hCA-II and hCA-IX in solution: the benzenesulfonamide hybrid showed double-digit nM inhibitory constant (K_i_^app^) towards both hCA-IX and hCA-II, and the AAZ hybrid showed single-digit nM K_i_^app^ towards both hCA-II and hCA-IX. (**Figure 4D, Supplementary Figure S15**). These results are somewhat different from the yeast surface titration assays, in which the benzenesulfonamide hybrid exhibited single-digit nM binding affinity to hCA-IX and a double-digit nM affinity to hCA-II. The AAZ hybrid’s strong inhibitory effect (single-digit nM) differs from the yeast-based triple-digit nM K_D_ to hCA-IX and high double-digit nM K_D_ to hCA-II. This discrepancy could be due to confounding factors in the yeast display experiments that lead to a lack of a plateau of target binding during titration experiments as discussed in *Survey and characterization of hybrids using flow cytometry*. Consistent with prior work, AAZ is such a potent pharmacophore that **4** alone exhibits potencies that appear to be indistinguishable from the potencies of clicked and thiol-maleimide hybrids. In the case of benzenesulfonamide, hybrids appear to exhibit stronger inhibitory effects than small molecule **1** alone, where the rate of product formation with target incubated in the presence of 50 nM hybrid is comparable to rate of product formation with target incubated with 200 nM small molecule **1** alone. This suggests that the integration of benzenesulfonamide into the Fn structure yields a synergistic hybrid structure in which the ncAA-containing Fn positions the pharmacophore such that the inhibitory potency is increased in the hybrid in comparison to either component alone. These inhibitory assays confirm that ncAA-containing hybrids retain inhibitory activity, confirming that assays performed on the yeast surface can be used to identify functional hybrids. However, it is also important to note that factors such as the multivalency of yeast display (and antigens) may confound quantitative comparisons. Overall, solution-based assays validate and extend the notion that “hybrid space” contains numerous functional, inhibitory molecules that can be identified with high-throughput experimentation strategies.

### Conclusions

In this work, we varied chemical elements of previously reported Fn-PEG_n_-AAZ hybrids to investigate a combinatorial set of Fn-sulfonamide hybrids constructed using ncAAs and click chemistry. To efficiently broaden the scope of these investigations, we also established a 96-well plate-based CuAAC protocol that greatly increases throughput of hybrid construction and evaluation. We identified several new functional hybrids, including benzenesulfonamide hybrids that exhibit potencies greater than the potency of the benzenesulfonamide warhead alone. These discoveries are striking in the context of the substantial number of chemical differences between the click hybrids and the “parent” thiol-maleimide hybrids as well as the moderate breadth of molecular variations evaluated; further chemical explorations of “hybrid space” are expected to yield additional functional hybrids with intriguing properties. While the yeast display format enabled efficient construction and evaluation of hybrids, we found that some caution is needed to ensure that data arising from assays on the yeast surface is not overinterpreted. While flow cytometry characterization of yeast displayed hybrids is high-throughput, weak binding signals can be lost over extended sample collection times. Yeast surface titration can yield rapid affinity characterizations of the clones, but in some cases we observed that the resulting K_D_ values do not necessarily predict the inhibitory properties of the clones in solution, possibly due to avidity via multimerization of the antigen and secondary reagents on the yeast surface as well as the potential for passive (non-inhibitory) binding. Future studies can take these limitations into account, use directly labeled antigens (to reduce multimerization), and include solution-based affinity characterizations of the most promising clones to ensure that binding and inhibitory behaviors are rigorously characterized.

The results described in this study demonstrate the potential to use clickable ncAAs to generate chemically diverse protein-small molecule hybrids. Stop codon suppression machinery enables precise control over the positions and chemical structures of ncAAs within binding protein scaffolds, and bioorthogonal functional groups facilitate efficient conjugations using workflows that are readily implemented without the use of specialized chemistry instrumentation^21, 22, 30, 31^. The CuAAC reactions used here also decrease the potential for nonspecific yeast surface modifications in comparison to thiol modifications (free thiols are known to be present on the surface of *S. cerevisiae*)^32, 33^. Synergistic combinations of small molecule warhead, linker structure, and binding protein needed to produce potent, specific inhibitors present a set of challenging design problems for which few precedents are available. High-throughput experimentation on the yeast surface enables the rapid discovery of inhibitors from large collections while simultaneously yielding data to better inform future hybrid design efforts. The ncAA-mediated production of clickable conjugates described here complements and extends prior work from our group to “chemically enhance” binding proteins by introducing photocrosslinkable and proximity-induced crosslinkable groups in binding proteins^23, 24^. This growing toolkit and numerous related strategies combining proteins and small molecules^34–37^ provide exciting opportunities to combinatorially evaluate hybrids for properties not accessible in either proteins or small molecules alone.

## References

1. Oprea, T. I.; Bologa, C. G.; Brunak, S.; Campbell, A.; Gan, G. N.; Gaulton, A.; Gomez, S. M.; Guha, R.; Hersey, A.; Holmes, J.; Jadhav, A.; Jensen, L. J.; Johnson, G. L.; Karlson, A.; Leach, A. R.; Ma’ayan, A.; Malovannaya, A.; Mani, S.; Mathias, S. L.; McManus, M. T.; Meehan, T. F.; von Mering, C.; Muthas, D.; Nguyen, D.-T.; Overington, J. P.; Papadatos, G.; Qin, J.; Reich, C.; Roth, B. L.; Schürer, S. C.; Simeonov, A.; Sklar, L. A.; Southall, N.; Tomita, S.; Tudose, I.; Ursu, O.; Vidović, D.; Waller, A.; Westergaard, D.; Yang, J. J.; Zahoránszky-Köhalmi, G., Unexplored therapeutic opportunities in the human genome. Nature Reviews Drug Discovery 2018, 17 (5), 317–332.

2. Kasperkiewicz, P.; Poreba, M.; Groborz, K.; Drag, M., Emerging challenges in the design of selective substrates, inhibitors and activity-based probes for indistinguishable proteases. FEBS J 2017, 284 (10), 1518–1539.

3. Wulff, H.; Christophersen, P.; Colussi, P.; Chandy, K. G.; Yarov-Yarovoy, V., Antibodies and venom peptides: new modalities for ion channels. Nat Rev Drug Discov 2019, 18 (5), 339–357.

4. Ayoub, M. A.; Crepieux, P.; Koglin, M.; Parmentier, M.; Pin, J. P.; Poupon, A.; Reiter, E.; Smit, M.; Steyaert, J.; Watier, H.; Wilkinson, T., Antibodies targeting G protein-coupled receptors: Recent advances and therapeutic challenges. MAbs 2017, 9 (5), 735–741.

5. Hudson, B. D.; Smith, N. J.; Milligan, G., Experimental challenges to targeting poorly characterized GPCRs: uncovering the therapeutic potential for free fatty acid receptors. Adv Pharmacol 2011, 62, 175–218.

6. Jo, M.; Jung, S. T., Engineering therapeutic antibodies targeting G-protein-coupled receptors. Exp Mol Med 2016, 48, e207.

7. Topiol, S., Current and Future Challenges in GPCR Drug Discovery. Methods Mol Biol 2018, 1705, 1–21.

8. Devy, L.; Dransfield, D. T., New Strategies for the Next Generation of Matrix-Metalloproteinase Inhibitors: Selectively Targeting Membrane-Anchored MMPs with Therapeutic Antibodies. Biochem Res Int 2011, 2011, 191670.

9. Devy, L.; Huang, L.; Naa, L.; Yanamandra, N.; Pieters, H.; Frans, N.; Chang, E.; Tao, Q.; Vanhove, M.; Lejeune, A.; van Gool, R.; Sexton, D. J.; Kuang, G.; Rank, D.; Hogan, S.; Pazmany, C.; Ma, Y. L.; Schoonbroodt, S.; Nixon, A. E.; Ladner, R. C.; Hoet, R.; Henderikx, P.; Tenhoor, C.; Rabbani, S. A.; Valentino, M. L.; Wood, C. R.; Dransfield, D. T., Selective inhibition of matrix metalloproteinase-14 blocks tumor growth, invasion, and angiogenesis. Cancer Res 2009, 69 (4), 1517–26.

10. Dufour, A.; Overall, C. M., Missing the target: matrix metalloproteinase antitargets in inflammation and cancer. Trends Pharmacol Sci 2013, 34 (4), 233–42.

11. Sexton, D. J.; Chen, T.; Martik, D.; Kuzmic, P.; Kuang, G.; Chen, J.; Nixon, A. E.; Zuraw, B. L.; Forteza, R. M.; Abraham, W. M.; Wood, C. R., Specific inhibition of tissue kallikrein 1 with a human monoclonal antibody reveals a potential role in airway diseases. Biochem J 2009, 422 (2), 383–92.

12. Zhang, J.; Valianou, M.; Simmons, H.; Robinson, M. K.; Lee, H. O.; Mullins, S. R.; Marasco, W. A.; Adams, G. P.; Weiner, L. M.; Cheng, J. D., Identification of inhibitory scFv antibodies targeting fibroblast activation protein utilizing phage display functional screens. FASEB J 2013, 27 (2), 581–9.

13. Prassas, I.; Eissa, A.; Poda, G.; Diamandis, E. P., Unleashing the therapeutic potential of human kallikrein-related serine proteases. Nature Reviews Drug Discovery 2015, 14, 183.

14. Chen, J.; Sawyer, N.; Regan, L., Protein-protein interactions: general trends in the relationship between binding affinity and interfacial buried surface area. Protein Sci 2013, 22 (4), 510–5.

15. Miller, D. W.; Dill, K. A., Ligand binding to proteins: the binding landscape model. Protein Sci 1997, 6 (10), 2166–79.

16. Schneider, T. L.; Mathew, R. S.; Rice, K. P.; Tamaki, K.; Wood, J. L.; Schepartz, A., Increasing the kinase specificity of k252a by protein surface recognition. Org Lett 2005, 7 (9), 1695–8.

17. Gower, C. M.; Thomas, J. R.; Harrington, E.; Murphy, J.; Chang, M. E.; Cornella-Taracido, I.; Jain, R. K.; Schirle, M.; Maly, D. J., Conversion of a Single Polypharmacological Agent into Selective Bivalent Inhibitors of Intracellular Kinase Activity. ACS Chem Biol 2016, 11 (1), 121–31.

18. Andersson, T.; Lundquist, M.; Dolphin, G. T.; Enander, K.; Jonsson, B.-H.; Nilsson, J. W.; Baltzer, L., The Binding of Human Carbonic Anhydrase II by Functionalized Folded Polypeptide Receptors. Chemistry & Biology 2005, 12 (11), 1245–1252.

19. Yang, J.; Koruza, K.; Fisher, Z.; Knecht, W.; Baltzer, L., Improved molecular recognition of Carbonic Anhydrase IX by polypeptide conjugation to acetazolamide. Bioorg Med Chem 2017, 25 (20), 5838–5848.

20. Lewis, A. K.; Harthorn, A.; Johnson, S. M.; Lobb, R. R.; Hackel, B. J., Engineered protein-small molecule conjugates empower selective enzyme inhibition. Cell Chem Biol 2021.

21. Stieglitz, J. T.; Kehoe, H. P.; Lei, M.; Van Deventer, J. A., A Robust and Quantitative Reporter System To Evaluate Noncanonical Amino Acid Incorporation in Yeast. ACS Synth Biol 2018, 7 (9), 2256–2269.

22. Stieglitz, J. T.; Van Deventer, J. A., High-Throughput Aminoacyl-tRNA Synthetase Engineering for Genetic Code Expansion in Yeast. ACS Synth Biol 2022, 11 (7), 2284–2299.

23. Alcala-Torano, R.; Islam, M.; Cika, J.; Lam, K. H.; Jin, R.; Ichtchenko, K.; Shoemaker, C.; Van Deventer, J., Yeast Display Enables Identification of Covalent Single Domain Antibodies Against Botulinum Neurotoxin Light Chain A. 2022.

24. Islam, M.; Kehoe, H. P.; Lissoos, J. B.; Huang, M.; Ghadban, C. E.; Berumen Sanchez, G.; Lane, H. Z.; Van Deventer, J. A., Chemical Diversification of Simple Synthetic Antibodies. ACS Chem Biol 2021, 16 (2), 344–359.

25. El-Gazzar, M. G.; Nafie, N. H.; Nocentini, A.; Ghorab, M. M.; Heiba, H. I.; Supuran, C. T., Carbonic anhydrase inhibition with a series of novel benzenesulfonamide-triazole conjugates. J Enzyme Inhib Med Chem 2018, 33 (1), 1565–1574.

26. Maren, T. H.; Conroy, C. W., A new class of carbonic anhydrase inhibitor. Journal of Biological Chemistry 1993, 268 (35), 26233–26239.

27. Bonardi, A.; Nocentini, A.; Bua, S.; Combs, J.; Lomelino, C.; Andring, J.; Lucarini, L.; Sgambellone, S.; Masini, E.; McKenna, R.; Gratteri, P.; Supuran, C. T., Sulfonamide Inhibitors of Human Carbonic Anhydrases Designed through a Three-Tails Approach: Improving Ligand/Isoform Matching and Selectivity of Action. J Med Chem 2020, 63 (13), 7422–7444.

28. Alterio, V.; Hilvo, M.; Di Fiore, A.; Supuran, C. T.; Pan, P.; Parkkila, S.; Scaloni, A.; Pastorek, J.; Pastorekova, S.; Pedone, C.; Scozzafava, A.; Monti, S. M.; De Simone, G., Crystal structure of the catalytic domain of the tumor-associated human carbonic anhydrase IX. Proc Natl Acad Sci U S A 2009, 106 (38), 16233–8.

29. Van Deventer, J. A.; Kelly, R. L.; Rajan, S.; Wittrup, K. D.; Sidhu, S. S., A switchable yeast display/secretion system. Protein Eng Des Sel 2015, 28 (10), 317–25.

30. Mocharla, V. P.; Colasson, B.; Lee, L. V.; Röper, S.; Sharpless, K. B.; Wong, C.-H.; Kolb, H. C., In Situ Click Chemistry: Enzyme-Generated Inhibitors of Carbonic Anhydrase II. Angewandte Chemie 2005, 117 (1), 118–122.

31. Manetsch, R.; Krasinski, A.; Radic, Z.; Raushel, J.; Taylor, P.; Sharpless, K. B.; Kolb, H. C., In situ click chemistry: enzyme inhibitors made to their own specifications. J Am Chem Soc 2004, 126 (40), 12809–18.

32. Lorenzen, I.; Eble, J. A.; Hanschmann, E. M., Thiol switches in membrane proteins - Extracellular redox regulation in cell biology. Biol Chem 2021, 402 (3), 253–269.

33. Brandes, N.; Reichmann, D.; Tienson, H.; Leichert, L. I.; Jakob, U., Using quantitative redox proteomics to dissect the yeast redoxome. J Biol Chem 2011, 286 (48), 41893–41903.

34. Kim, C. H.; Axup, J. Y.; Lawson, B. R.; Yun, H.; Tardif, V.; Choi, S. H.; Zhou, Q.; Dubrovska, A.; Biroc, S. L.; Marsden, R.; Pinstaff, J.; Smider, V. V.; Schultz, P. G., Bispecific small molecule-antibody conjugate targeting prostate cancer. Proc Natl Acad Sci U S A 2013, 110 (44), 17796–801.

35. Rogers, O. C.; Rosen, D. M.; Antony, L.; Harper, H. M.; Das, D.; Yang, X.; Minn, I.; Mease, R. C.; Pomper, M. G.; Denmeade, S. R., Targeted delivery of cytotoxic proteins to prostate cancer via conjugation to small molecule urea-based PSMA inhibitors. Sci Rep 2021, 11 (1), 14925.

36. Chiang, M. J.; Holbert, M. A.; Kalin, J. H.; Ahn, Y. H.; Giddens, J.; Amin, M. N.; Taylor, M. S.; Collins, S. L.; Chan-Li, Y.; Waickman, A.; Hsiao, P. Y.; Bolduc, D.; Leahy, D. J.; Horton, M. R.; Wang, L. X.; Powell, J. D.; Cole, P. A., An Fc domain protein-small molecule conjugate as an enhanced immunomodulator. J Am Chem Soc 2014, 136 (9), 3370–3.

37. Liu, F.; Ul Amin, T.; Liang, D.; Park, M. S.; Alhamadsheh, M. M., Enhancing the Pharmacokinetic Profile of Interleukin 2 through Site-Specific Conjugation to a Selective Small-Molecule Transthyretin Ligand. J Med Chem 2021, 64 (19), 14876–14886.

