## Supplementary data for "Systematic evaluation of protein-small molecule hybrids on the yeast surface"

### Supplementary figures

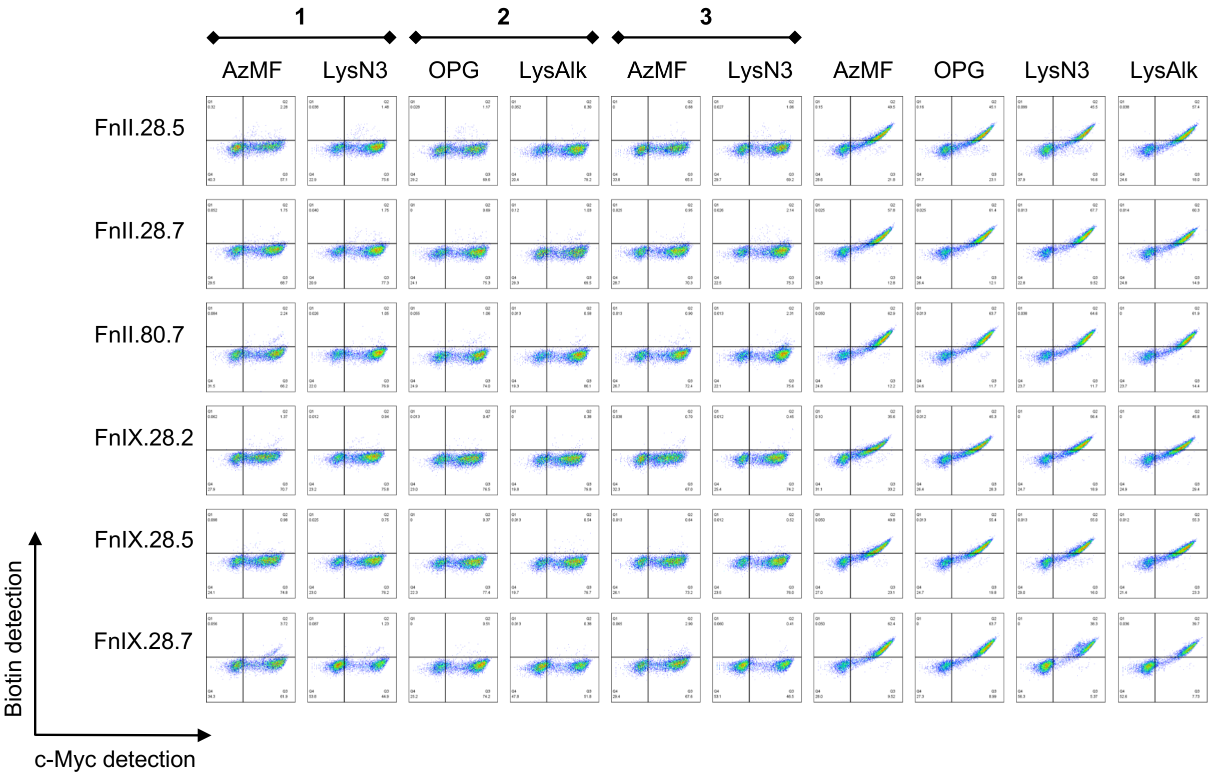


**Figure S1** Biotin detection level of the Fn-sulfonamide (**1, 2, 3**) hybrids and the ncAA-containing Fns after 2-step click chemistry. After the first-step small molecule click chemistry, the hybrids have few clickable ncAAs left to react with the clickable biotin probes in the second step, therefore showing very low biotin detection. Meanwhile the Fns not reacted with small molecules have all their clickable ncAA for the biotin probes to click with, therefore showing high biotin levels. 2-step biotin click is performed after every click chemistry conjugation and the data are consistent with the data shown here.

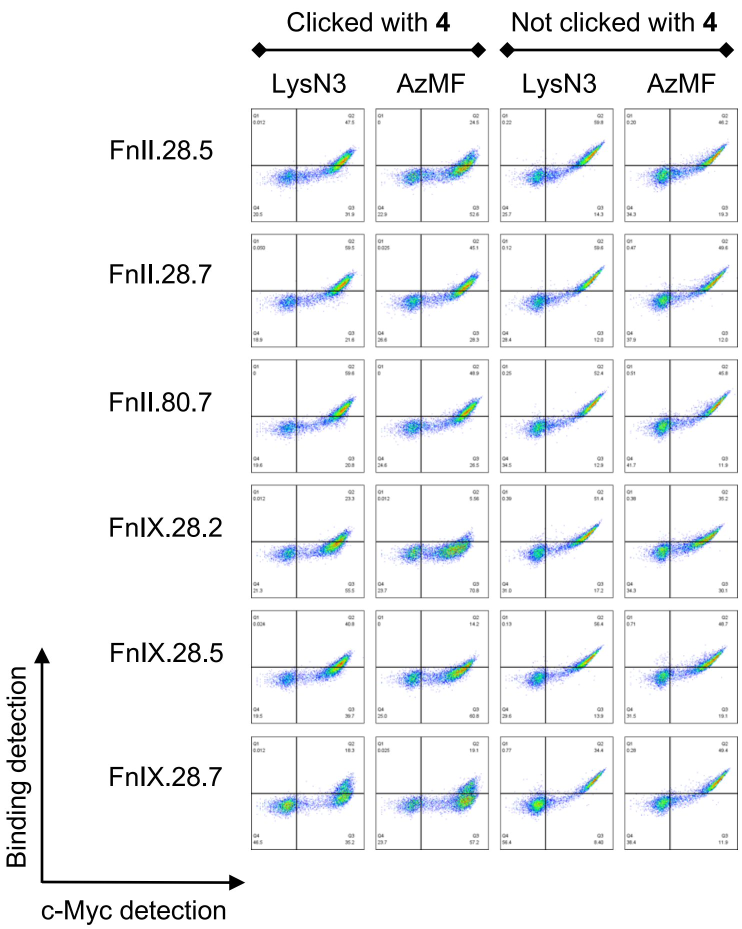


**Figure S2** Biotin detection level of the Fn-sulfonamide (**1, 2, 3**) hybrids and the ncAA-containing Fns after 2-step click chemistry. After the first-step small molecule click chemistry, the hybrids have few clickable ncAAs left to react with the clickable biotin probes in the second step, therefore showing very low biotin detection. Meanwhile the Fns not reacted with small molecules have all their clickable ncAA for the biotin probes to click with, therefore showing high biotin levels. 2-step biotin click is performed after every click chemistry conjugation and the data are consistent with the data shown here.

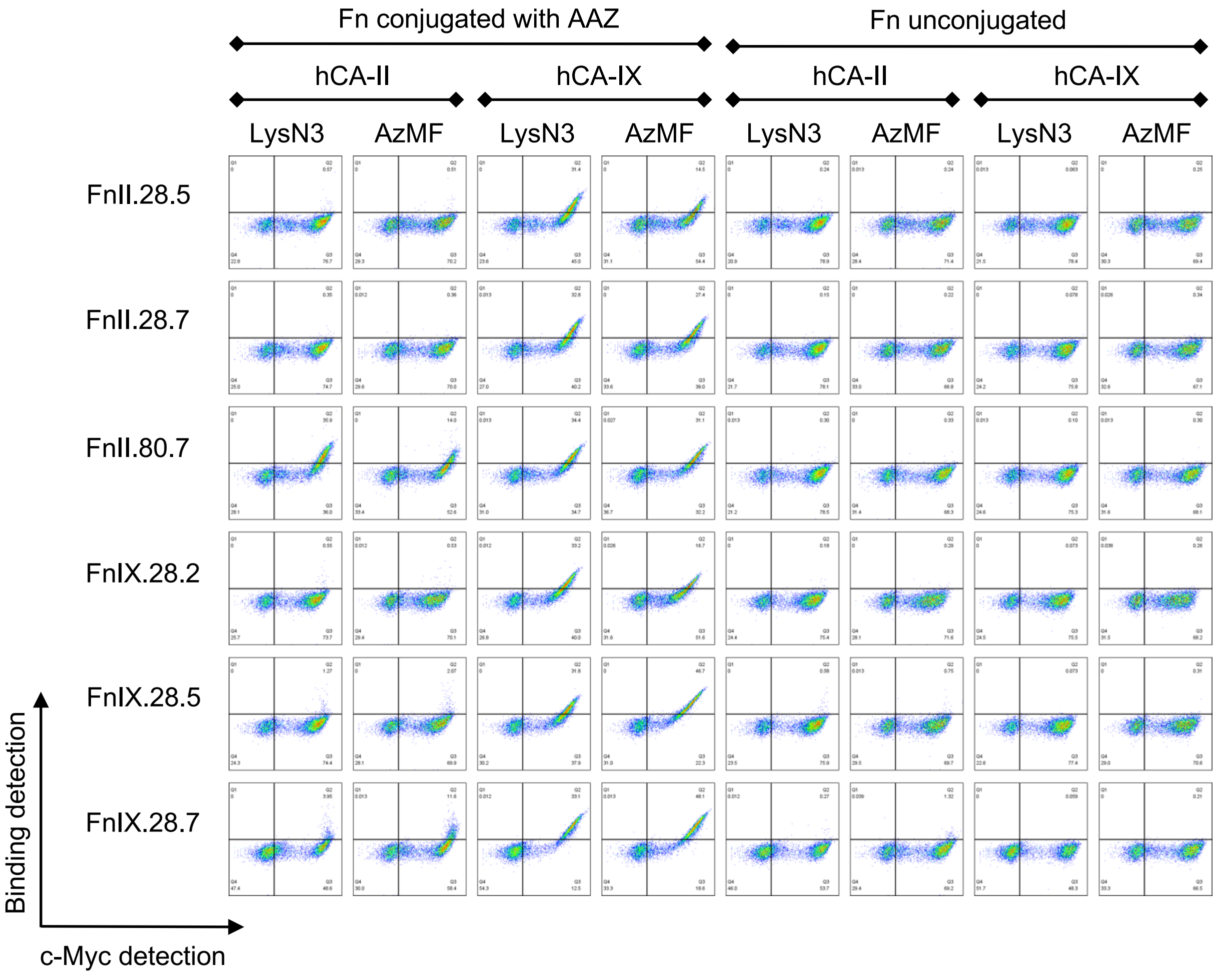


**Figure S3** Flow cytometry data for the Fn-AAZ yeast surface 50 nM hCA binding screening. Corresponding to Figure 2B. All the dot plots are shown in logarithmic scales.

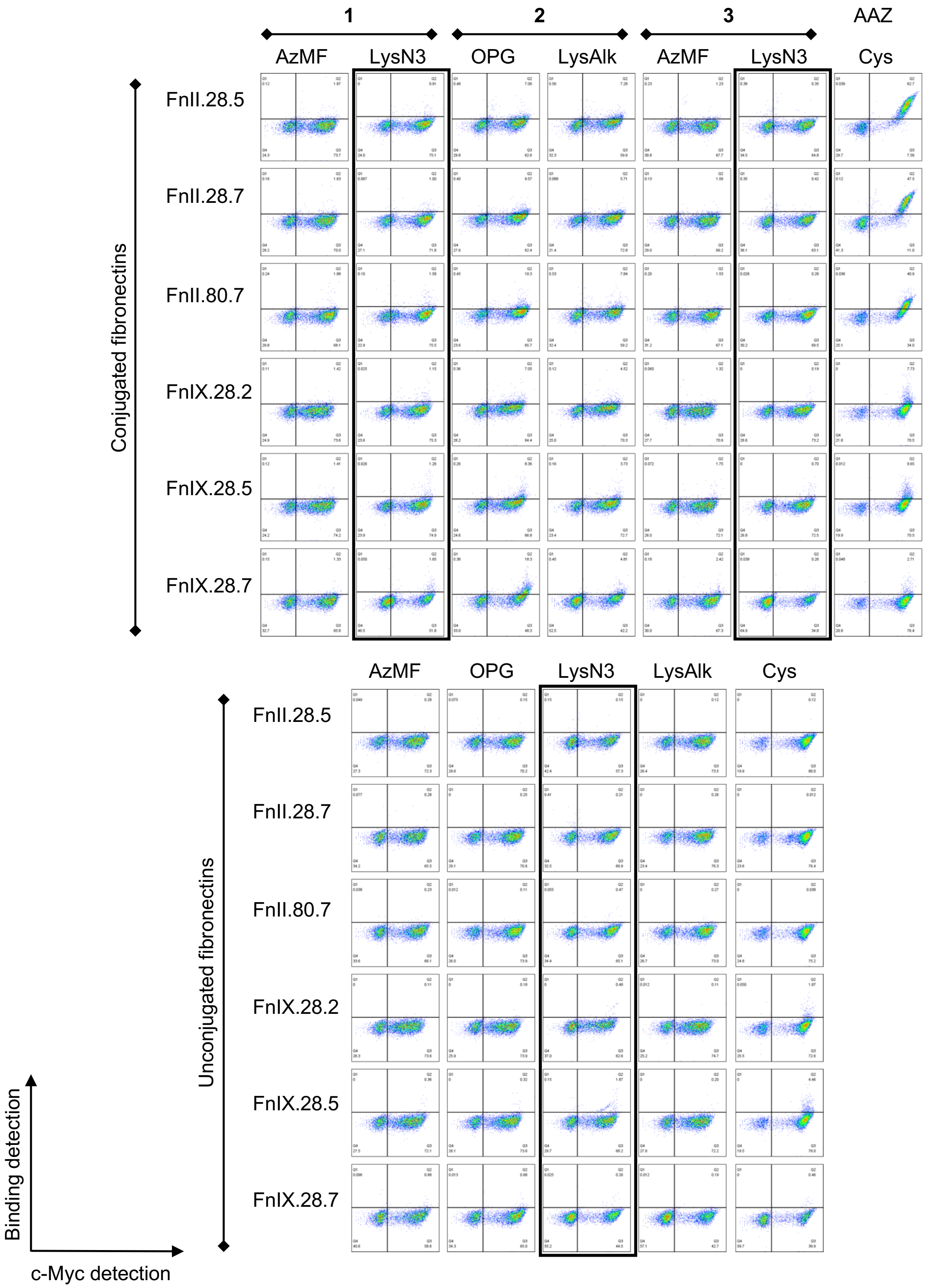


**Figure S4** Flow cytometry data of the yeast surface 50 nM hCA-II binding screening for Fns conjugated with other sulfonamides (**1**, **2**, **3**). Corresponding to Figure 2C. Data in black rectangles are taken in a later experiment under the same binding and labeling conditions.

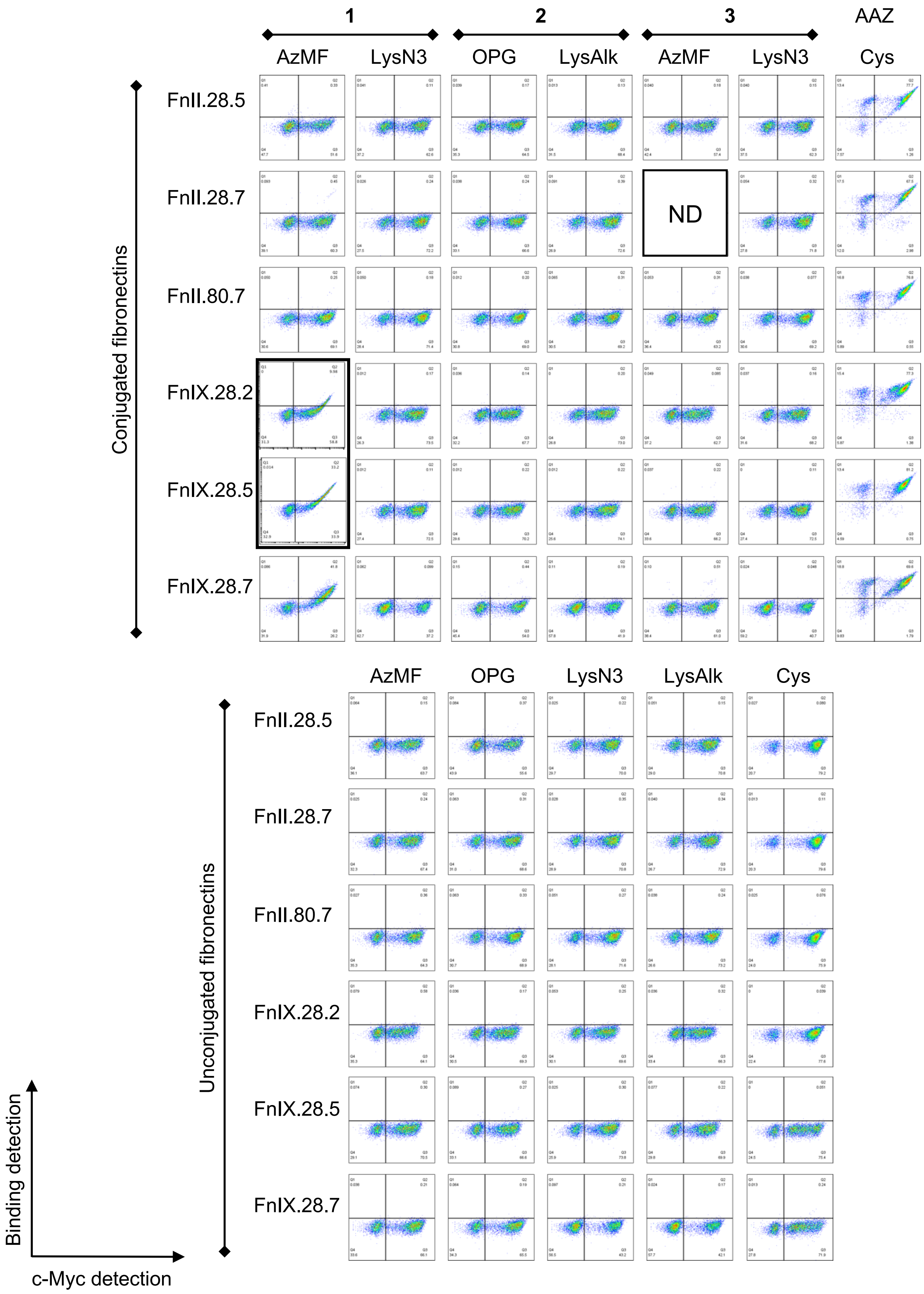
1
2
3
AAZ
AzMF
LysN3
OPG
LysAlk
AzMF
LysN3
Cys
Conjugated fibronectins
FnII.28.5
FnII.28.7
FnII.80.7
FnIX.28.2
FnIX.28.5
FnIX.28.7
ND
AzMF
OPG
LysN3
LysAlk
Cys
Unconjugated fibronectins
FnII.28.5
FnII.28.7
FnII.80.7
FnIX.28.2
FnIX.28.5
FnIX.28.7
Binding detection
c-Myc detection

**Figure S5** Flow cytometry data of the yeast surface 50 nM hCA-IX binding screening for Fns conjugated with other sulfonamides (**1**, **2**, **3**). Corresponding to Figure 2C. Data in black rectangle are taken in a previous experiment under the same binding and labeling conditions. One sample was not collected due to technical issues and is labeled ND (not determined).

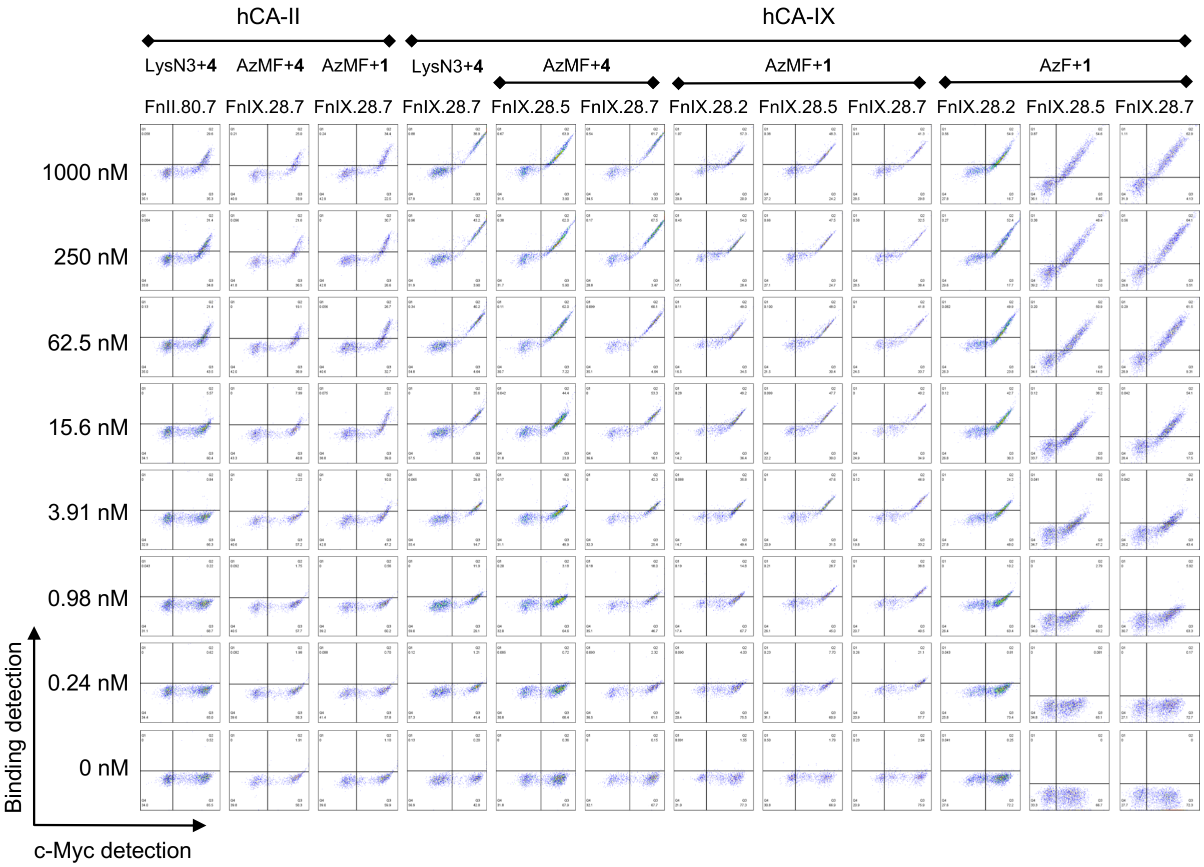


**Figure S6** Flow cytometry dot plots of binding titrations used to determine binding affinities (corresponding to Table 1. Fns were incubated with different concentrations of hCA-**II** or hCA-IX (starting from 1 μM with seven subsequent 4-fold dilutions and a final, 0 nM hCA concentration) and analyzed for binding. Labeling conditions are listed in Table S4. Each condition represented here was evaluated in technical triplicates. Samples were all prepared under the same conditions. Data depicted in the two rightmost columns was obtained on a different instrument (BioRad s3e) than the other data depicted in this figure (Attune NxT) due to technical difficulties with the Attune NxT on the day of analysis.

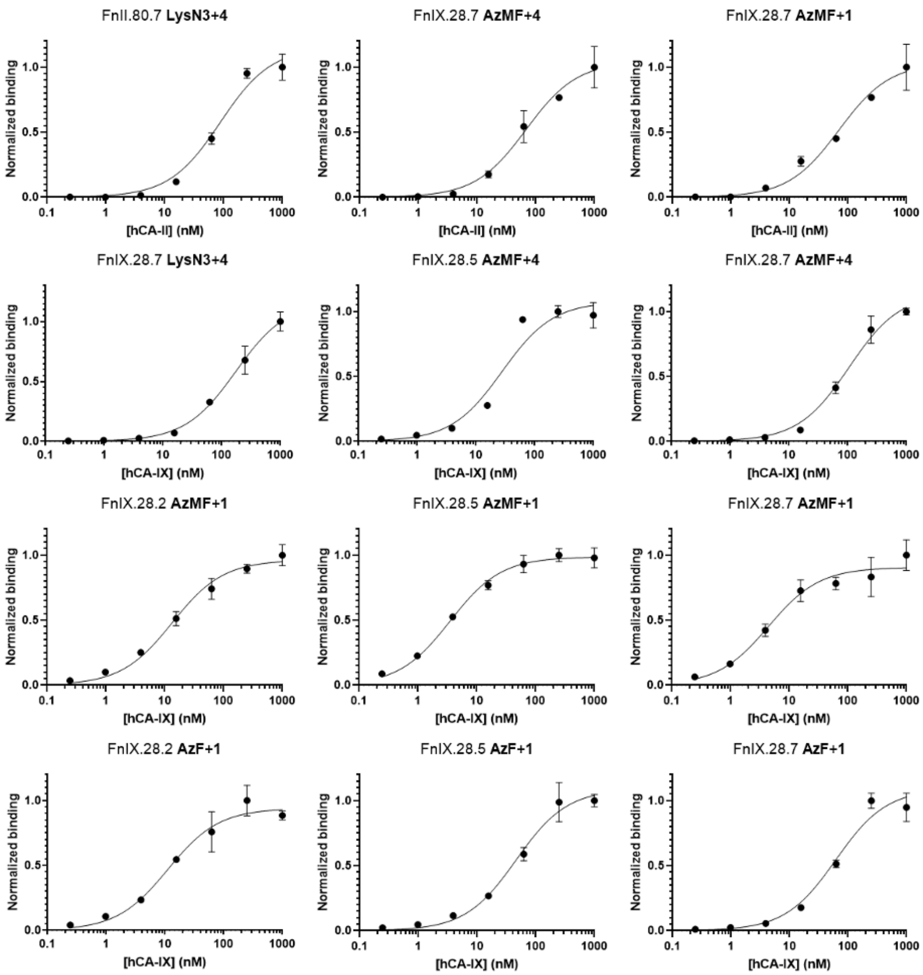


**Figure S7** Titration curves for ncAA-containing Fn-small molecules hybrids studied in this work. Binding assays were performed with different concentrations of biotinylated IgG (starting from 1 μM with subsequent 4-fold dilutions) for each Fn-small molecule hybrid (see also Supplementary Figure S7). Median Fluorescence Intensity (MFI) data for hCA detection was corrected for background detection by subtracting MFI of non-displaying populations from the MFI of displaying populations for each sample. Background subtracted MFI data was then normalized to the highest MFI detection for each hybrid (obtained with 1 μM antigen), using GraphPad Prism 9, and plotted as a function of hCA concentration. Each condition was tested in technical triplicates with average values and standard error reported here. Binding affinities ($K_D$) were calculated using the “Receptor binding – Saturation binding” model and “One site -- Specific binding” equation in GraphPad Prism 9; these results are reported in **Table 1**.

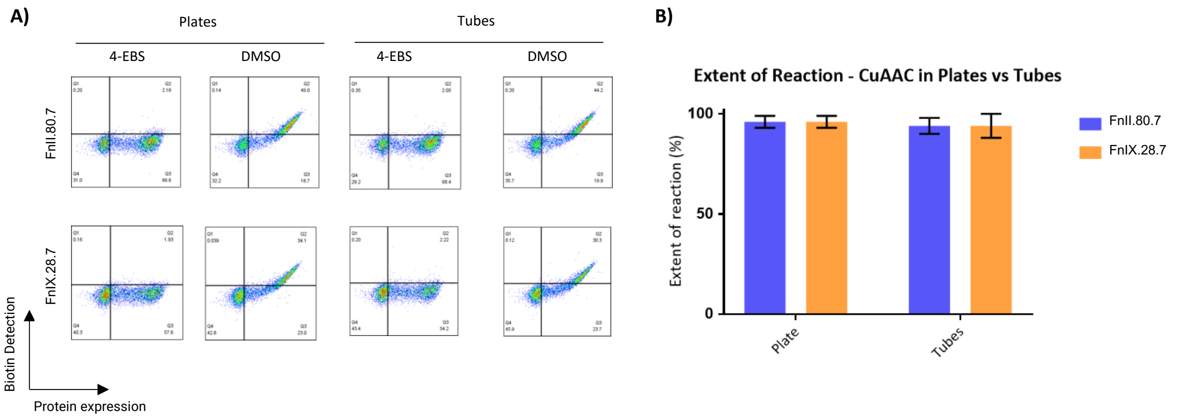


**Figure S8** Comparison of CuAAC between AzMF and 4-ethynylbenzenesulfonamide (4-EBS) in plates versus tubes. **A)** Representative flow cytometry plots of biotin CuAAC on cells displaying FnII.80.7 or FnIX.28.7, performed in plates or in tubes as indicated. A first round of CuAAC was performed with 4-ethynylbenzenesulfonamide (4-EBS) or with DMSO as a control. **B)** Extent of reaction of FnII.80.7 and FnIX.28.7 in plates versus tubes. Error bars indicate standard deviation for technical triplicates.

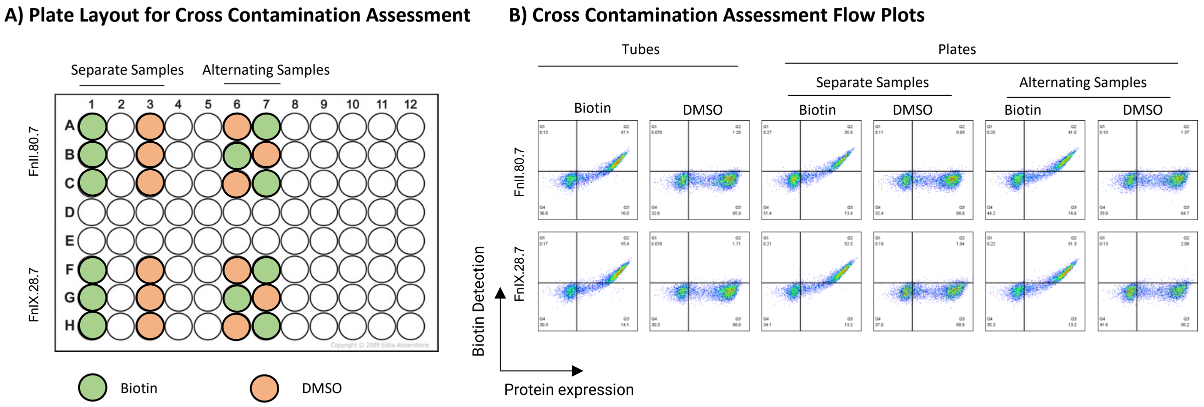


**Figure S9** Assessment of cross contamination in CuAAC in plates. A) Plate layout for cross-contamination assessment experiment. Samples reacted with biotin are represented in green, and samples reacted with DMSO are represented in orange. B) Representative flow plots of biotin CuAAC showcasing the lack of cross contamination when performing CuAAC in plates.

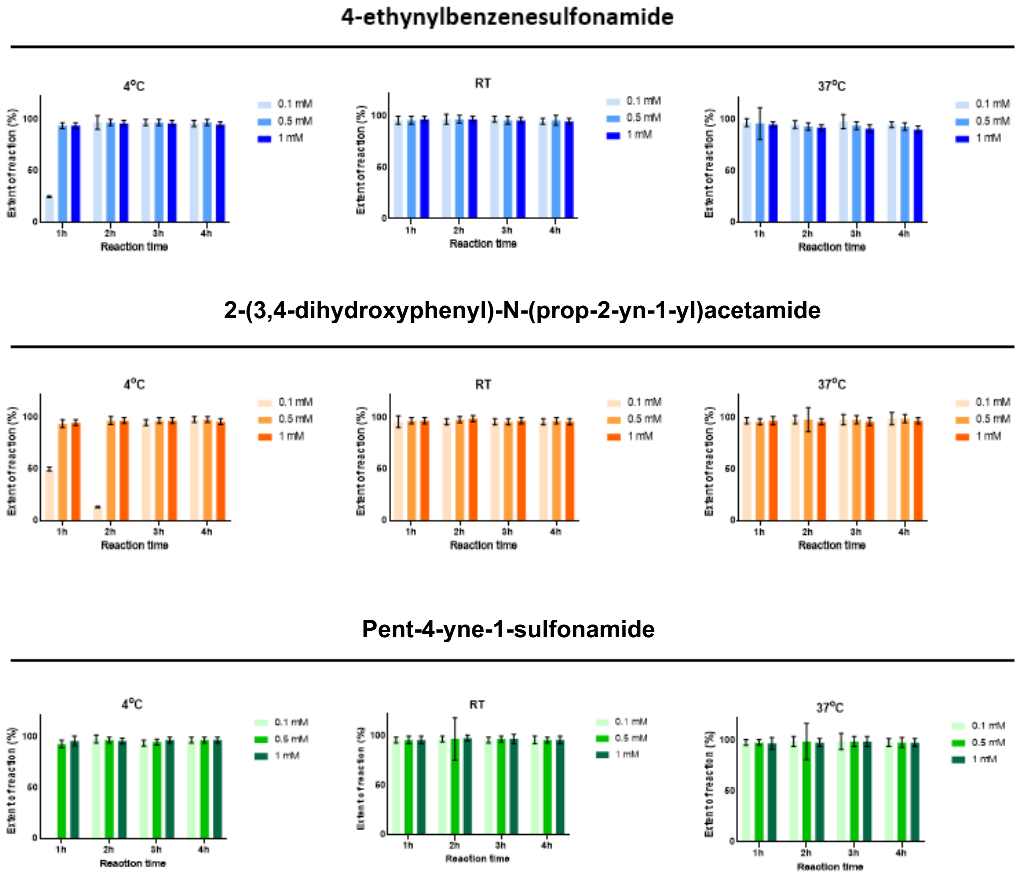


**Figure S10** Extent of reaction for each condition studied under CuAAC screening. Similarity across conditions reflects flexibility of reaction conditions, except for low temperature and low small molecule concentration conditions.

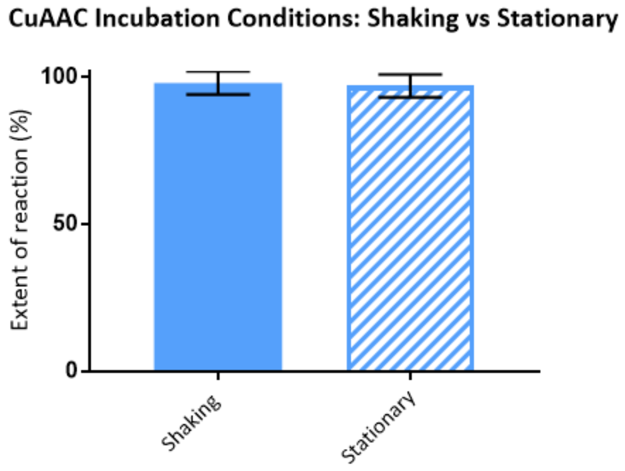


**Figure S11** Average extent of reaction values from triplicate samples comparing shaking versus stationary incubation conditions. Extent of reaction is not affected by shaking or stationary conditions for this specific conjugation between FnII.80.7 and 4-ethynylbenzenesulfonamide.

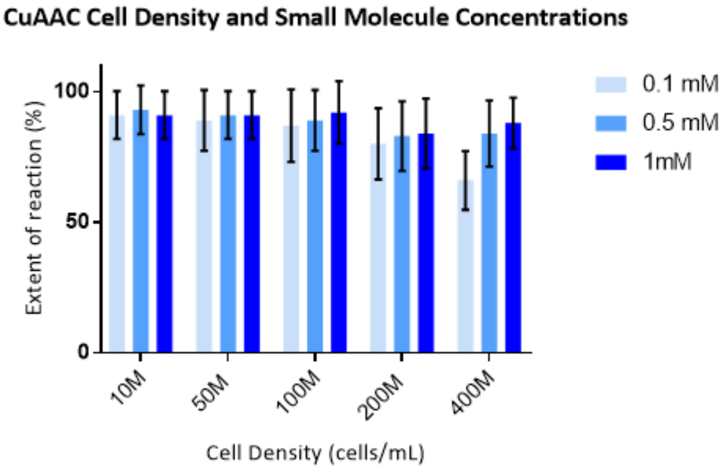


**Figure S12** Extent of reaction of 4-ethynylbenzenesulfonamide with FnII.80.7 under different cell density conditions and small molecule concentrations.

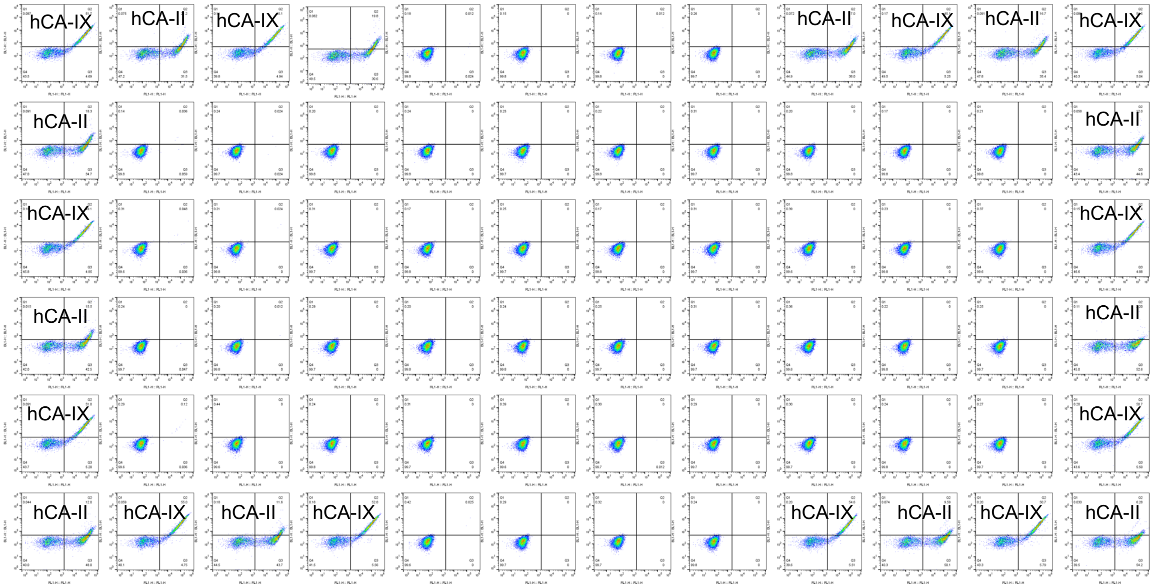


| hCA-II | | | | | | | | | | | | |
|---|---|---|---|---|---|---|---|---|---|---|---|---|
| | 1 | 2 | 3 | 4 | 5 | 6 | 7 | 8 | 9 | 10 | 11 | 12 |
| B | | 267 | | 266 | | | | | 190 | | 193 | |
| C | 201 | | | | | | | | | | | 137 |
| D | | | | | | | | | | | | |
| E | 147 | | | | | | | | | | | 80 |
| F | | | | | | | | | | | | |
| G | 125 | | 131 | | | | | | | 112 | | 95 |

| hCA-IX | | | | | | | | | | | | |
|---|---|---|---|---|---|---|---|---|---|---|---|---|
| | 1 | 2 | 3 | 4 | 5 | 6 | 7 | 8 | 9 | 10 | 11 | 12 |
| B | 3760 | | 3708 | | | | | | | 3166 | | 2821 |
| C | | | | | | | | | | | | |
| D | 3569 | | | | | | | | | | | 2899 |
| E | | | | | | | | | | | | |
| F | 3394 | | | | | | | | | | | 2657 |
| G | | 3325 | | 3161 | | | | | 2906 | | 2448 | |

**Figure S13**. Signal decay of hCA binding detection on a 12 by 6 plate layout. In each well there are 1 million yeast cells displaying FnIX.28.7 with AzMF and small molecule **1.** Samples on the edge of the plate are incubated with 50 nM hCA-II or hCA-IX and then labeled for flow cytometry. All 96 wells are resuspended together after labeling and data are collected horizontally at a speed of around 1 min per well. The specific binding MFI are listed in a heat map format with arbitrary color scales.

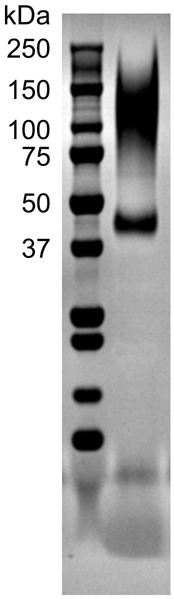


**Figure S14** SDS-PAGE analysis of Protein A-purified FnIX.28.7 induced with AzMF. Gels were loaded under reducing conditions and stained with coomassie SimplyBlue SafeStain.

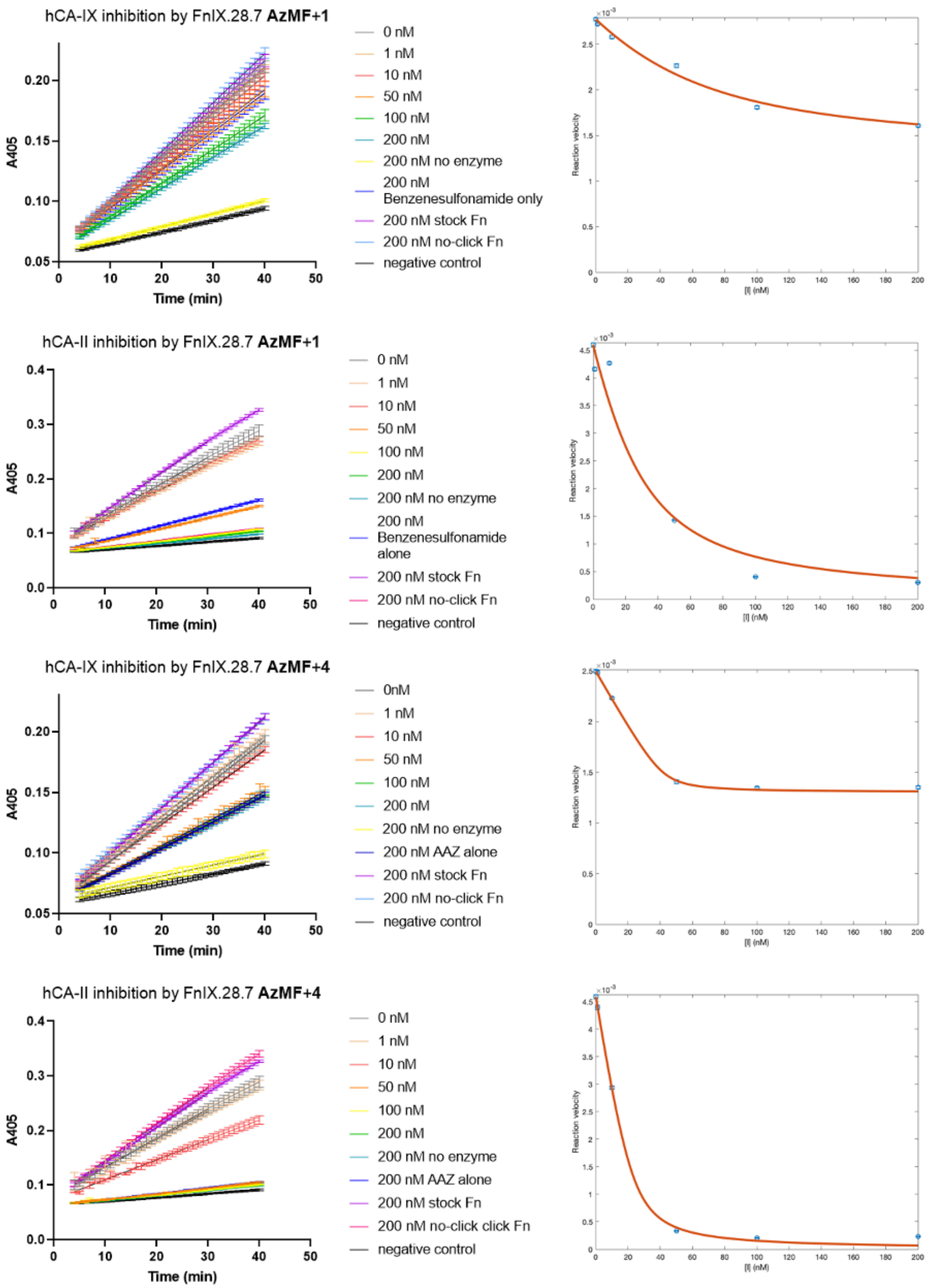


**Figure S15**. Experiment data of hCA inhibition assays using 4-NPA and fitted curve using the tight binding Morrison model. Error bars are ±1 standard error. In the case of hCA-IX inhibition, the glycine in the enzyme formulation hydrolyzed 4-NPA and caused a higher baseline of enzyme reactivity (**Figure S16**). Therefore, we set 0.0013 as an observed baseline for hCA9 assays from the AAZ hybrid inhibition data and fitted the model with respect to this baseline activity.

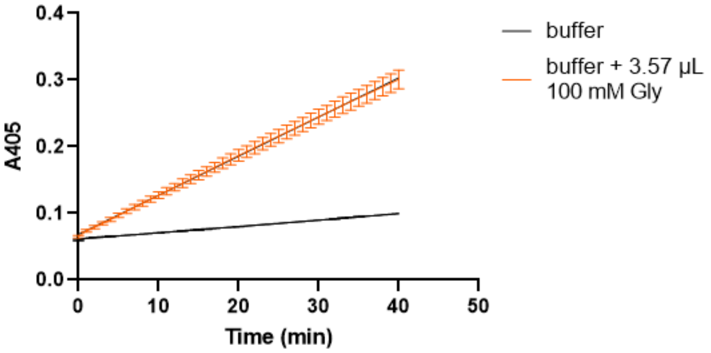


**Figure S16**. 4-NPA is hydrolyzed by the glycine in the hCA-IX formulation. 100 mM glycine in water solution was added to the buffer and 4-NPA mixture to match the amount of glycine added alongside hCA-IX in the inhibition assays and hydrolyzation of the product was observed.

### Supplementary Tables

**Table S1.** Extent of reaction for sulfonamide (**1**-**3**) hybrids in the screening

| | **AzMF+1** | **LysN3+1** | **OPG+2** | **LysAlk+2** | **AzMF+3** | **LysN3+AS** |
|---|---|---|---|---|---|---|
| FnII.28.5 | 94% ± 14% | 98% ± 7% | 98% ± 12% | 99% ± 6% | 96% ± 8% | 98% ± 7% |
| FnII.28.7 | 98% ± 5% | 98% ± 4% | 98% ± 5% | 98% ± 7% | 98% ± 7% | 98% ± 6% |
| FnII.80.7 | 98% ± 6% | 98% ± 4% | 99% ± 5% | 98% ± 5% | 98% ± 6% | 98% ± 4% |
| FnIX.28.2 | 94% ± 18% | 97% ± 7% | 97% ± 9% | 97% ± 11% | 95% ± 15% | 97% ± 11% |
| FnIX.28.5 | 96% ± 10% | 98% ± 6% | 98% ± 7% | 98% ± 13% | 98% ± 8% | 97% ± 5% |
| FnIX.28.7 | 97% ± 4% | 99% ± 3% | 98% ± 3% | 99% ± 3% | 97% ± 3% | 99% ± 3% |

**Table S2.** Extent of reaction for AAZ (**4**) hybrids in the screening

| | **AzMF+4** | **LysN3+4** |
|---|---|---|
| FnII.28.5 | 71% ± 6% | 56% ± 4% |
| FnII.28.7 | 52% ± 5% | 47% ± 4% |
| FnII.80.7 | 58% ± 3% | 50% ± 3% |
| FnIX.28.2 | 81% ± 11% | 68% ± 6% |
| FnIX.28.5 | 74% ± 7% | 54% ± 5% |
| FnIX.28.7 | 91% ± 4% | 86% ± 3% |

**Table S3**. DNA sequence of the 6 fibronectins used in this work. The codon for cysteine is highlighted in cyan and is switched to TAG codon to allow ncAA incorporation. The NheI and BamHI restriction sites introduced before and after the framework to allow cloning into a pCT40 vector are highlighted in yellow. XmaI site is used instead of BamHI site for pCHA-FcSup-TAA vector.

| Name of the clone | Short Code | DNA sequence |
|---|---|---|
| FnII.28.5 | 2.2 | GCTAGCTCCTCCGACTCTCCGCGTAACCTGGAGGTTACCA<br>ACGCAACTCCGAACTCTCTGACTATTTCTTGGGACTATCCT<br>GATTGCGCTATCTATTACCGTATCACCTACGGCGAAACTG<br>GTGGTAACTCCCCGAGCCAGGAATTCACTGTTCCGGGAAC<br>TAATACTAATGCGACCATCAGCGGTCTGAAACCGGGCCAG<br>GATTATACCATTACCGTGTACGCTGTAACTGATTATAGCCT<br>TAGTAATTCAAACCCAATCAGCATCAATTATCGCACCGAAA<br>TCGACAAACCGTCTCAGGGATCC |
| FnII.28.7 | 2.3 | GCTAGCTCCTCCGACTCTCCGCGTAACCTGGAGGTTACCA<br>ACGCAACTCCGAACTCTCTGACTATTTCTTGGGACGATTCT<br>GCGGCTTGCGCTCTCTATTACCGTATCACCTACGGCGAAA<br>CTGGTGGTAACTCCCCGAGCCAGGAATTCACTGTTCCGGG<br>AACTTCTTATAATGCGACCATCAGCGGTCTGAAACCGGGC<br>CAGGATTATACCATTACCGTGTACGCTGTAGCTGATTATAG<br>CCTTCAGTATTCAAACCCAATCAGCATCAATTATCGCACCG<br>AAATCGACAAACCGTCTCAGGGATCC |
| FnII.80.7 | 2.14 | GCTAGCTCCTCCGACTCTCCGCGTAACCTGGAGGTTACCA<br>ACGCAACTCCGAACTCTCTGACTATTTCTTGGGACTATCAT<br>TATCTTGCTCTCTATTACCGTATCACCTACGGCGAAACTGG<br>TGGTAACTCCCCGAGCCAGGAATTCACTGTTCCGGGAAAT<br>ACTTATGCGACCATCAGCGGTCTGAAACCGGGCCAGGATT<br>ATACCATTACCGTGTACGCTGTAACTGATTATGACTGCATT<br>TCAAACCCAATCAGCATCAATTATCGCACCGAAATCGACAA<br>ACCGTCTCAGGGATCC |
| FnIX.28.2 | 9.1 | GCTAGCTCCTCCGACTCTCCGCGTAACCTGGAGGTTACCA<br>ACGCAACTCCGAACTCTCTGACTATTTCTTGGGACTATCAT<br>CAGAATGGTTGCGCTGTTTCTTACCGTATCACCTACGGCG<br>AAACTGGTGGTAACTCCCCGAGCCAGGAATTCACTGTTCC<br>GGGTTATTACGATACCTATTCCGCGACCATCAGCGGTCTG<br>AAACCGGGCCAGGATTATACCATTACCGTGTACGCTGTAA<br>CCGGTTATAACGACGATTCAAACCCAATCAGCATCAATTAT<br>CGCACCGAAATCGACAAACCGTCTCAGGGATCC |
| FnIX.28.5 | 9.5 | GCTAGCTCCTCCGACTCTCCGCGTAACCTGGAGGTTACCA<br>ACGCAACTCCGAACTCTCTGACTATTTCTTGGGACGATTAT<br>CAGAATTGCGCTGTTTCTTACCGTATCACCTACGGCGAAAC<br>TGGTGGTAACTCCCCGAGCCAGGAATTCACTGTTCCGGGT<br>TATTACGATACCTATTCCGCGACCATCAGCGGTCTGAAACC<br>GGGCCAGGATTATACCATTACCGTGTACGCTGTAACCGGT |

| | | |
|---|---|---|
| | | TATAACGACGATTCAAACCCAATCAGCATCAATTATCGCAC CGAAATCGACAAACCGTCTCAGGGATCC |
| FnIX.28.7 | 9.7 | GCTAGCTCCTCCGACTCTCCGCGTAACCTGGAGGTTACCA ACGCAACTCCGAACTCTCTGACTATTTCTTGGGACGATTAT TTGTTGTGCGTTTTTTCTTACCGTATCACCTACGGCGAAAC TGGTGGTAACTCCCCGAGCCAGGAATTCACTGTTCCGGGA TATACTAATACTGTGACCATCAGCGGTCTGAAACCGGGCC AGGATTATACCATTACCGTGTACGCTGTAACTACTTATGAC CTTGATTCAAACCCAATCAGCATCAATTATCGCACCGAAAT CGACAAACCGTCTCAGGGATCC |

**Table S4**. Primers used for cloning. The TAG codon in the position of the TGC codon are highlighted in red.

| | | |
|---|---|---|
| | pCT 40-FN_fwd | GGTGGGTCTGGTGGGGGCGGATCTGCTAGCTC CTCCGACTCTCCG |
| | pCT 40-FN_rev | CAAGTCCTCTTCAGAAATAAGCTTTTGTTCGGAT CCCTGAGACGGTTTGTCGATTTCGGT |
| FnII.28.5 | FN2-2TAG_f | TGACTATTTCTTGGGACTATCCTGATTAGGCTAT CTATTACCGTATCACCTACGGCGAA |
| | FN2-2TAG_r | AGGTGATACGGTAATAGATAGCCTAATCAGGATA GTCCCAAGAAATAGTCAGAGAGTTCG |
| FnII.28.7 | FN2-3TAG_f | GACTATTTCTTGGGACGATTCTGCGGCTTAGGCT CTCTATTACCGTATCACCTACGGCG |
| | FN2-3TAG_r | CGCCGTAGGTGATACGGTAATAGAGAGCCTAAG CCGCAGAATCGTCCCAAGAAATAGTC |
| FnII.80.7 | FN2-14TAG_f | GTGTACGCTGTAACTGATTATGACTAGATTTCAA ACCCAATCAGCATCAATTATCGCA |
| | FN2-14TAG_r | AATTGATGCTGATTGGGTTTGAAATCTAGTCATA ATCAGTTACAGCGTACACGGTAATGG |
| FnIX.28.2 | FN9-1 TAG_f | TATTTCTTGGGACTATCATCAGAATGGTTAGGCT GTTTCTTACCGTATCACCTACGGCG |
| | FN9-1 TAG_r | TGATACGGTAAGAAACAGCCTAACCATTCTGATG ATAGTCCCAAGAAATAGTCAGAGAG |
| FnIX.28.5 | FN9-5 TAG_f | GACTATTTCTTGGGACGATTATCAGAATTAGGCT GTTTCTTACCGTATCACCTACGGCG |
| | FN9-5 TAG_r | GTGATACGGTAAGAAACAGCCTAATTCTGATAAT CGTCCCAAGAAATAGTCAGAGAGTTC |
| FnIX.28.7 | FN9-7 TAG_f | ACTATTTCTTGGGACGATTATTTGTTGTAGGTTTT TTCTTACCGTATCACCTACGGCGAA |
| | FN9-7 TAG_r | TGATACGGTAAGAAAAAACCTACAACAAATAATC GTCCCAAGAAATAGTCAGAGAGTTCG |
| | pCHA -FcSup-TAA -FN-Fwd | CCATACGACGTTCCAGACTACGCTGCTAGCTCC TCCGACTCTCCGCGTAACCT |
| | pCHA -FcSup-TAA -FN-Rev | TTTGTCGGAACTTTTAGGTTCTACCCCGGGCTGA GACGGTTTGTCGATTTCGGTGCG |

**Table S5** List of plasmids for Fn and suppression machineries

| | Backbone | Auxotrophic marker | Clone | Incorporates ncAA |
|---|---|---|---|---|
| Display plasmid | pCT40 | Trp | FnII.28.5 | N/A |
| | | | FnII.28.7 | |
| | | | FnII.80.7 | |
| | | | FnIX.28.2 | |
| | | | FnIX.28.5 | |
| | | | FnIX.28.7 | |
| Secretion plasmid | pCHA-FcSup-TAA | Trp | FnIX.28.2 | |
| | | | FnIX.28.5 | |
| | | | FnIX.28.7 | |
| TAG codon suppression machinery | pRS315 | Leu | AcFRS | *p*-azido-L-phenylalanine (AzF) |
| | | | LeuOmeRS | *p*-azidomethyl-L-phenylalanine (AzMF) |
| | | | | *p*-propargyloxyphenylalanine (OPG) |
| | | | B-LysAlkRS-3 (Kan marker) | *N*-ε-((2-Azidoethoxy)carbonyl)- L-lysine (LysN3) |
| | | | | *N*-ε-propargyloxycarbonyl-L-lysine (LysAlk) |

**Table S6** Labeling antibody used for flow cytometry and western blotting

| Target | Primary labeling | Secondary labeling |
|---|---|---|
| c-Myc tag | Chicken anti c-Myc (1:250) | Goat anti-Chicken Alexa Fluor 647 (1:500) |
| hCA-II (His tag) | Mouse anti His (3D5) (1:250) | Goat anti-Mouse Alexa Fluor 488 (1:500) |
| hCA-IX (human Fc tag) | N/A | Goat anti-Human IgG Fc Alexa Fluor 488 (1:125) |
| biotin | N/A | Streptavidin Alexa Fluor 488 (1:500) |

**Table S7** Small molecules used in the plate-based high-throughput hybrid construction

| | Chemical Name | ncAA |
|---|---|---|
| 5 | CUDC-101 | AzMF |
| 6 | 10-undecynoic acid | AzMF |
| 7 | 6-heptynoic acid | AzMF |
| 8 | N-hydroxy-10-undecinamide | AzMF |
| 9 | 2-(3,4-dihydroxyphenyl)-N-(prop-2-yn-1-yl)acetamide | AzMF |
| 1 | 4-Ethynylbenzene-1-sulfonamide | AzMF |
| 3 | Pent-4-yne-sulfonamide | AzMF |
| 4 | N-(5-sulfamoyl-1,3,4-thiadiazol-2-yl)hex-5-ynamide | AzMF |
| 2 | 4-azidobenzenesulfonamide | OPG |
| 10 | 2-(2-Propyn-1-yloxy)ethanesulfonyl fluoride | AzMF |
| 11 | But-3-yne-1-sulfonyl fluoride | AzMF |
| 12 | 4-ethynylbenzene-1-sulfonyl fluoride | AzMF |
| 13 | N-(hept-6-yn-1-yl)-2,3-dihydroxybenzamide | AzMF |
| 14 | N-[2-(3,4-dihydroxyphenyl)ethyl]hept-6-ynamide | AzMF |
| 15 | 3-(but-3-yn-1-yl)-1-[2-(3,4-dihydroxyphenyl)ethyl]thiourea | AzMF |
| 16 | N-(2-azidoethyl)-[2,2'-bipyridine]-5-carboxamide | OPG |
| 17 | 4-azido-N-({[2,2'-bipyridine]-6-yl}methyl)butanamide | OPG |
| 18 | N-(5-azidopentyl)-[2,2'-bipyridine]-5-carboxamide | OPG |
| 19 | N-(5-azidopentyl)-2-(pyridin-2-yl)quinoline-4-carboxamide | OPG |
| 20 | 2,3-diamino-5-bromo-N-(hept-6-yn-1-yl)benzamide | AzMF |
| 21 | 2-propynylurea | AzMF |
| 22 | 1-phenyl-3-prop-2-yn-1-ylurea | AzMF |
| 23 | 4-(Dihydroxyborophenyl)acetylene | AzMF |
| 24 | 4-(propargylaminocarbonyl)phenylboronic acid | AzMF |

**Table S8** Conditions for the hCA inhibition assay

| V (µL) | [FnIX.28.7-Fc] (nM) | [4-NPA] (mM) | [hCA] (nM) | Note |
|---|---|---|---|---|
| 100 | 0 | 2 | 100 | |
| | 1 | | 100 | |
| | 10 | | 100 | |
| | 50 | | 100 | |
| | 100 | | 100 | |
| | 200 | | 100 | |
| | 200 no-click Fn | | 100 | Fn mixed with small molecule and all click chemistry reagents but Cu. |
| | 200 nM stock Fn | | 100 | Fn not subjected to any CuAAC regents |
| | 200 no enzyme | | 0 | Control for hydrolyzation of 4-NPA from the hybrids |
| | 0 | | 0 | Negative control |
| | 200 nM small molecule alone | | 100 | |

### Materials and methods

*Introducing TAG codon into fibronectin DNA sequences*

Original fibronectin sequences[1] were cloned into pCT40 using the NheI and BamHI restriction sites. Primers encoding a TAG codon at the cysteine position were used to introduce the mutations to the fibronectins. Two DNA segments were amplified via PCR for each fibronectin to introduce the mutation at the desired position. Gel electrophoresis was performed on the PCR products to extract DNA fragments of the expected insert size, and then recombined with a pCT40 vector digested at the NheI and BamHI restriction sites using Gibson Assembly. The assemblies were transformed into chemically competent *E. coli* DH5αZ1, plated on LB plates containing ampicillin and grown at 37 °C overnight. Resulting colonies were picked and grown to saturation in 5 mL LB liquid media containing ampicillin. Plasmid DNA was then isolated from these cultures using GenCatch Plasmid DNA Mini-Prep kits. The cloned plasmids were digested at the NheI and BamHI sites to confirm the presence of correctly sized inserts by performing gel electrophoresis. Finally, the plasmids were sequence verified from Quintara Biosciences.

*Yeast display and stop codon suppression*

The yeast surface display of ncAA-containing protein was performed following previously established protocols[2]. For display constructs encoding proteins containing only canonical amino acids, display plasmids (pCTCON2 backbone, TRP marker) were transformed into Zymo competent RJY100 cells, plated on SD-CAA (–TRP –URA) solid media and allowed to grow for 2–3 days at 30 °C, until colonies appeared. For display constructs encoding a TAG codon, display constructs (pCTCON2 backbone, TRP marker) were transformed alongside aaRS/tRNA suppressor plasmids (pRS-315 plasmids, LEU marker, AcFRS for AzF, LeuOmeRS for AzMF and OPG, B-LysAlkRS-3 for LysN3 and LysAlk) using Zymo competent RJY100 cells, plated on SD-SCAA TRP –LEU –URA solid media and allowed to grow for 2–3 days at 30 °C, until colonies appeared. The details of display plasmid and aaRS/tRNA plasmid combinations used in this study are listed in Supplementary Table S3. For propagation, colonies were inoculated in 2–5 mL of the respective liquid media supplemented with penicillin-streptomycin at 1:100 dilution (+ Pen/Strep) and left to grow at 30 °C with shaking at 250–300 RPM until saturation (2–3 days). Saturated cultures were stored at 4 °C for future propagation. To propagate, 100–500 µL of cells from the saturated culture were pelleted and resuspended to an OD600 of 0.5–1 in 5 mL of fresh liquid media + Pen/Strep and grown overnight at 30 °C with shaking at 250–300 RPM. Once saturated, cultures were diluted to an OD600 of 1 and allowed to grow to mid-log phase (OD600 = 2–5; 4–8 hours) and then induced by pelleting and resuspending in the corresponding SG-SCAA media + Pen/Strep to an OD of 1. SG-CAA was used for cultures containing display plasmids only, SGSCAA –TRP –LEU –URA was used for cultures containing display plasmids as well as aaRS/tRNA suppression machinery. To facilitate ncAA incorporation, induction media for cultures with TAG-mutated display plasmids were supplemented with ncAAs to a final concentration of 1 mM of the L-isomer. Cells were grown in the induction media at 20 °C with shaking at 250–300 RPM for 15–18 hours before further experimentation.

*Production and purification of soluble Fn-Fc*

The Fn sequences were cloned into a previously established yeast secretion plasmid, pCHA-FcSup-TAA plasmid[3, 4] using NheI and XmaI restriction sites and Gibson Assembly. Isolation of PCR products of approximately the correct size and successful Gibson Assembly reactions, as well as general transformation and plasmid isolation steps were performed as described in the "Introduction of TAG codons into fibronectin DNA sequences" section. All plasmids were sequence verified with sequencing performed at Quintara Biosciences. Zymo competent RJY100 yeast cells were transformed with the pCHA-FcSup-TAA plasmids (TRP marker) containing the Fn sequences as well as pRS-315 plasmids (LEU marker) containing the amber codon suppression machinery (aaRS/tRNA pair, AcFRS for AzF, LeuOmeRS for AzMF and OPG, B-LysAlkRS-3 for LysN3 and LysAlk).

Fn-Fcs were secreted and purified according to previously established protocols[4] with the following modifications. Yeast culture was expanded in SDSCAA-Trp-Leu-Ura media at 30°C and then induced in YPG with 1 mM ncAA and 0.1% w/v bovine serum albumin (Proliant Biologicals) at 20°C, shaken at 275-300 rpm. Penicillin-Streptomycin is used to prevent contaminations. After four days of incubation, the culture was spun down for 35 minutes at 3214 rcf and the supernatant was buffered with 10x PBS, pH 7.4 and sterile filtered with 0.2 μM filter cups. The filtrate was passed twice through a protein A column, following which the Fn-Fc containing resin was washed three times with 10 mL PBS. The Fn-Fcs were then eluted using 7 mL 100 mM glycine, pH 3.0 and immediately neutralized with 0.7 mL 1 M Tris, pH 8.5. The eluant was buffer exchanged using Amicon Ultra-15 centrifugal filter units (30 kDa molecular weight cut-off, Millipore Sigma) into PBS, pH 7.4 and concentrated. Protein concentrations were measured via the absorbance of 280 nm light on a Nano Drop One instrument (Thermo Fisher). Protein purity was determined via sodium dodecyl sulfate polyacrylamide gel electrophoresis (SDS-PAGE) without the use of PNGase F (**Supplementary Figure S14**).

*Yeast surface and protein click chemistry*

Copper-catalyzed azide-alkyne cycloaddition (CuAAC) reactions on yeast surface were performed as described previously with some modifications specific to the small molecules used.[5, 6] Reaction container is 1.7 mL microcentrifuge tubes; reaction buffer is 220 μL 1× PBS pH 7.4. For cell surface reactions 2 M cells were used per reaction, reagents are 1) DMSO stock solution of small molecule or biotin probe (amount varies for different small molecules), 2) 3.8 μL $CuSO_4$/THPTA (1:2 ratio of 20 mM $CuSO_4$:50 mM THPTA), 3) 12.5 μL 100 mM aminoguanidine and 4) 12.5 μL 100 mM sodium ascorbate (in the given order), vortexing briefly in between each addition. For small molecule **1**, **2**, and **3** the reaction was done with 1 mM small molecule (2.5 μL 100 mM stock solution) for 2 hours at room temperature. For small molecule **4** and clickable biotin probes the reaction was done with 0.1 mM small molecule (1.25 μL 20 mM stock solution) for 15 minutes at room temperature, since longer reaction time or concentration of **4** will lead to more complete modification but loss of specific binding function. After the reaction the cells were washed three times in PBSA to prepare for flow cytometry assays.

Click chemistry for soluble protein is done in the same conditions as the yeast surface reactions but using purified Fn-Fc proteins. After conjugating the small molecules, the protein is buffer exchanged to reach a $10^7$-fold dilution of the reaction solution to terminate the reaction and remove excess small molecules. Fn-Fcs that had undergone reactions without the copper catalyst and Fn-Fcs that had not been subjected to click chemistry reactions were used as controls. Western blots were used to verify the presence of clickable ncAA and in the TAG-mutated Fn-Fcs as well as the absence of clickable ncAA in the conjugated Fn-Fcs. SDSPAGE using 4–12% Bis-Tris gels was performed on all samples in duplicate gels. One gel was stained with Coomassie SimplyBlue SafeStain to confirm the presence of protein. Western blots were performed by transferring the second PAGE gel to a nitrocellulose membrane using an iBlot2 Dry Blotting System (Life Technologies). Transferred membranes were blocked with 5% w/v BSA in TBS + 0.1% v/v Tween20, following which they were probed with streptavidin Alexa Fluor 488 in a 1:1000 dilution for the presence of biotin (**Figure 4B**).

*Initial binding screening flow cytometry and data analysis*
Analytical flow cytometry was performed as described in previous work.[5] To prepare Fns for flow cytometry analysis, 2 million freshly induced cells of each sample were washed three times in PBSA (1× PBS, pH 7.4, with 0.1% w/v BSA) and then conjugated with alkyne-azide click chemistry or cysteine-maleimide chemistry. Cells are then labeled in 96 well V-bottom plates for flow cytometry. Human CA-II (Sino Biological, 10478-H08E) or CA-IX (Sino Biological, 10107-H02H) were incubated with the cells at room temperature for 1 hour on an orbital shaker at 150 RPM. Following binding, all subsequent steps were performed on ice or in a centrifuge chilled to 4 °C. Cells were diluted in ice-cold PBSA and washed three times, also with ice-cold PBSA. Labeling for His tag or human Fc tag and the following fluorophore labelling was performed on ice for 10 minutes in the dark with 3 washes in between. Samples were diluted in and washed twice with ice-cold PBSA before final resuspension in PBSA for flow cytometry. The labeling reagents and concentrations used for primary and secondary labelling are listed in **Supplementary Table S6**. Labeling reagents were prepared in PBSA. Flow cytometry was performed on an Attune NxT flow cytometer (Life Technologies) in the Tufts University Science and Technology Center, and data was processed using FlowJo™ software. 10,000 events were collected per sample on the flow cytometer. One factor affecting the detected fluorescence level is the data collection time for a large number of samples. When we evaluated hCA binding as a function of well location in a 6 by 12 layout used in the binding survey, we noted that hybrids show decreasing binding signals as a function of plate position (**Supplementary Figure S13**). This led to us initially missing a hybrid exhibiting binding activity (FnIX.28.7 with AzMF+**1** binding to hCA-II) until it was identified in the high-throughput screening where there was less waiting time between labeling and flow cytometry analysis (This issue can be avoided in future work by increasing data collection speeds in the flow cytometer).

The extent of reaction of the cell surface CuAAC is calculated based on the labeling for c-Myc and conjugated biotin on the cell surface. The equations are listed below (Equation 1-3):

$$Extent\ of\ SM\ click\ reaction = 1 - Extent\ of\ Biotin\ click\ reaction \quad \text{(Equation 1)}$$

$$Extent\ of\ Biotin\ click\ reaction = \frac{MFI\ sample - MFI\ background}{MFI\ positive\ control - MFI\ background} \quad \text{(Equation 2)}$$

$$Extent\ of\ SM\ click\ reaction = 1 - \frac{MFI\ sample - MFI\ background}{MFI\ positive\ control - MFI\ background} \quad \text{(Equation 3)}$$

The errors are calculated based of the CV values of the MFI values (Equation 4-7)

$$Numerator\ CV = \sqrt{CV\ of\ MFI\ sample^2 + CV\ of\ MFI\ background^2} \quad \text{(Equation 4)}$$

$$Denominator\ CV = \sqrt{CV\ of\ MFI\ control^2 + CV\ of\ MFI\ background^2} \quad \text{(Equation 5)}$$

$$CV\ of\ final\ value = \sqrt{\frac{Numerator\ CV}{Numerator\ value}^2 + \frac{Denominator\ CV}{Denominator\ value}^2} \quad \text{(Equation 6)}$$

$$Propagated\ Error = Final\ value(extent\ of\ Biotin\ reaction) * CV\ of\ final\ value \quad \text{(Equation 7)}$$

*Yeast surface titration flow cytometry*

2 million freshly induced RJY100 cells displaying Fns were conjugated with small molecules with click chemistry in 1.7 mL microcentrifuge tubes. After the conjugation, cells were pelleted, washed three times with 1× PBSA, and then resuspended in 1 mL PBSA. To prepare for binding titration, 15,000 cells were added to wells of 96-well V-bottom plates. Human CA-II or CA-IX were serial diluted starting at 1 µM with seven subsequent 4-fold dilutions and a final PBSA blank. The enzyme dilutions were added to the wells (ensuring >10x excessive enzyme) and incubated on the orbital shaker at 150 RPM at room temperature for 2 hours. Following labeling and flow cytometry data collection was performed as described in the “Initial binding screening flow cytometry and data analysis” above, with the exception that 3,000 events were collected per sample instead of the usual 10,000 events. All titrations were done in technical triplicates. We assessed one hybrid titration at a time to avoid long waiting time between labeling and detecting. We kept the cell number constant across the series of hCA concentrations, therefore the display level is controlled and normalized in the final data analysis. Data analysis to obtain binding affinities ($K_D$) were done using steps like those outlined in section “Initial binding screening flow cytometry and data analysis” above, with a few modifications. Samples were gated to isolate single cells and gated on c-Myc detection levels to isolate positive (induced) and negative (uninduced) populations. MFI data for hCA-II or hCA-IX detection of induced populations and uninduced populations was retrieved. Background correction was performed by subtracting hCA detection MFI values of uninduced cell populations from MFI values of corresponding induced populations. On GraphPad Prism, the background corrected data for hCA detection were normalized for binding (by converted MFI values to ratios between the highest

hCA concentration and all subsequent lower concentrations) and then, using the "Receptor binding – Saturation binding" model and "One site -- Specific binding" equation, the data was fitted to a curve and $K_D$ estimated including standard error and 95% confidence interval.

*High-throughput fibronectin hybrid generation in 96-well plate*

Cell pellets were resuspended in 175 µL 1X PBS at pH 7.4. Then, 2 µL of small molecule (or DMSO for control) were added to each well individually, for a final concentration of 1 mM. Using a multichannel pipette, the 3 µL $CuSO_4$/THPTA aliquots in a separate plate were transferred to each sample, followed by 10 µL of aminoguanadine and 10 µL of sodium ascorbate. Both aminoguanidine and sodium ascorbate reach a final concentration of 5 mM. All steps were performed promptly and pipetting up and down several times after the addition of each reagent to ensure proper mixing. The plate was covered with aluminum foil and left on an orbital shaker at 150 rpm at room temperature for 4 hours, except for the case of small molecule **4**, which is reacted for 15 min at 0.1 mM. Once the reaction time was over, cells were washed three times with ice-cold PBSA and then left as pellets which could be either used immediately after or saved at 4°C overnight for binding assays with hCA-II or hCA-IX, or for the second step of the reaction. For the biotin probe click chemistry, cell pellets were resuspended in 176 µL 1X PBS at pH 7.4. All reagents are the same except for the small molecule or DMSO, which were substituted by 20 mM biotin alkyne. To each well, 1 µL of biotin-alkyne was added individually. The rest of the procedure is the same as the first step described above, adding 3 µL of $CuSO_4$/THPTA, 10 µL of aminoguanidine and 10 µL of sodium ascorbate in that order. The plate was covered up with aluminum foil and on left on an orbital shaker at 150 rpm at room temperature for 15 minutes. Once the reaction was over, cells were washed three times in ice-cold PBSA and left as pellets to be prepared for labeling. For negative controls, the first reaction with small molecule or DMSO is omitted, and the cells are only subjected to this reaction with the biotin-alkyne probe. We tested different combinations of reaction conditions such as temperature, reagent concentration, and cell density, and found out that most reactions behave well in a wide range of reaction conditions (**Supplementary Figure S10-12**), facilitating high-throughput generation of hybrids of various compositions.

*Carbonic anhydrase inhibition assay*

Carbonic anhydrase esterase activity was measured by monitoring the hydrolyzation of 4-nitrophenyl acetate (4-NPA) to 4-nitrophenol and acetic acid in the presence of the enzyme. The increase of 4-nitrophenol increased absorbance at 405 nm, which is recorded with a SpectraMax i3x microplate reader. Human CA-II (hCA-II) (Sino Biological 10478-H08E) or Human CA-IX (hCA-IX) (Sino Biological 10107-H02H) was incubated with the inhibitor at 0, 1, 10, 50, 100, or 200 nM for 30 minutes, and then the substrate was added right before the start of the recording. The absorbance was measured every minute for 40 minutes at room temperature. Reaction buffer was 12.5 mMTris, 75 mM NaCl, pH 7.5. The 100 mM stock solution of 4-NPA in acetone was carefully diluted 5x in reaction buffer, and then 10 µL of the dilution was added to the reaction in a final enzymatic assay volume of 100 µL containing 2 mM 4-NPA.

Concentration of enzymes was fixed at 100 nM for hCA-II or hCA-IX. The inhibition assays were done in triplicates.

*Inhibition assay data analysis*

The absorbance at 405 nM over time (after the initial 4 minutes, data points that are higher than the two readings before and after it in the curve are removed as outliners) was linearly fit in GraphPad Prism. Once the reaction rates and the corresponding inhibitor concentrations are obtained, the rate of spontaneous hydrolyzation of 4-NPA without enzyme was subtracted from the enzymatic rates. Since we are using 4-nitrophenyl acetate (4-NPA) for the detection of hCA activity, it is hard to determine the actual concentration of 4-NPA dissolved in the solution as 4-NPA is not very soluble in aqueous buffers. The $K_m$ of the hCA-II and hCA-IX to 4-NPA is also hard to determine precisely but is estimated to be higher than the concentration of 4-NPA used in inhibition assays. Therefore we opted to determine the apparent inhibition constant $K_i^{app}$ of the hybrids following the Hackel lab's method.[1] When we tried to fit the Morrison equation for tight binding to our data, we encountered 2 problems: 1) The hCA-II and AAZ hybrid inhibition assay data could not fit well to the curve with the nominal enzyme concentration and the estimates for $K_i^{app}$ are unrealistically small to the point of negative values; and 2) our hCA-IX inhibition data exhibit a residue level of 4-NPA hydrolysis that is not inhibitable with our hybrid inhibitors or AAZ small molecule alone. Problem 1 suggests that the nominal enzyme concentration is an unrealistic overestimate of the actual active enzyme concentration in the solution. A similar overestimate of enzyme concentration leading to negative $K_i$ estimates was reported by Kuzmič et. al (2000), and to overcome this problem they determined the enzyme concentration and the $K_i^{app}$ together in the model fitting.[7] We decided to use the AAZ hybrids inhibition data to determine the enzyme concentration since AAZ hybrids are more inhibitory than the benzenesulfonamdie hybrids, therefore giving us a more reliable enzyme concentration that is not likely overestimated. The residue level of 4-NPA hydrolysis is due to the 100 mM glycine in the formulation of hCA-IX, as we observed glycine hydrolyzing 4-NPA on its own (**Supplementary Figure S16**). Therefore we adjusted the baseline of hCA-IX inhibition data to 0.0013 to account for the substrate hydrolysis from glycine. To obtain apparent $K_i$ the data was fitted to Equation 8 in MATLAB_2017a using *nlinfit*, solving for $K_i^{app}$ and enzyme concentration ($E_T$). Default options were used except for *TolX* and *TolFun*, which are set to 1e-15. Initial velocities $v_0$ were set to the observed values from the enzyme and substrate only conditions (0 nM inhibitors). The function *nlparci* was used to determine the 68% confidence interval (alpha=0.32)

$$\frac{v_i}{v_0} = 1 - \frac{(E_T + I_T + K_i^{app}) - \sqrt{(E_T + I_T + K_i^{app})^2 - 4E_T I_T}}{2E_T} \qquad \text{(Equation 8)}$$

### References

1. Lewis, A. K.; Harthorn, A.; Johnson, S. M.; Lobb, R. R.; Hackel, B. J., Engineered protein-small molecule conjugates empower selective enzyme inhibition. *Cell Chem Biol* **2021**.

2. Hershman, R. L.; Rezhdo, A.; Stieglitz, J. T.; Van Deventer, J. A., Engineering Proteins Containing Noncanonical Amino Acids on the Yeast Surface. In *Yeast Surface Display*, Traxlmayr, M. W., Ed. Springer US: New York, NY, 2022; pp 491-559.

3. Van Deventer, J. A.; Kelly, R. L.; Rajan, S.; Wittrup, K. D.; Sidhu, S. S., A switchable yeast display/secretion system. *Protein Eng Des Sel* **2015,** *28* (10), 317-25.

4. Islam, M.; Kehoe, H. P.; Lissoos, J. B.; Huang, M.; Ghadban, C. E.; Berumen Sanchez, G.; Lane, H. Z.; Van Deventer, J. A., Chemical Diversification of Simple Synthetic Antibodies. *ACS Chem Biol* **2021,** *16* (2), 344-359.

5. Stieglitz, J. T.; Kehoe, H. P.; Lei, M.; Van Deventer, J. A., A Robust and Quantitative Reporter System To Evaluate Noncanonical Amino Acid Incorporation in Yeast. *ACS Synth Biol* **2018,** *7* (9), 2256-2269.

6. Van Deventer, J. A.; Le, D. N.; Zhao, J.; Kehoe, H. P.; Kelly, R. L., A platform for constructing, evaluating, and screening bioconjugates on the yeast surface. *Protein Eng Des Sel* **2016,** *29* (11), 485-494.

7. Kuzmic, P.; Elrod, K. C.; Cregar, L. M.; Sideris, S.; Rai, R.; Janc, J. W., High-throughput screening of enzyme inhibitors: simultaneous determination of tight-binding inhibition constants and enzyme concentration. *Anal Biochem* **2000,** *286* (1), 45-50.
